# A diet-dependent host metabolite shapes the gut microbiota to protect from autoimmunity

**DOI:** 10.1101/2023.11.02.565382

**Authors:** Margaret Alexander, Vaibhav Upadhyay, Rachel Rock, Lorenzo Ramirez, Kai Trepka, Patrycja Puchalska, Diego Orellana, Qi Yan Ang, Caroline Whitty, Jessie A. Turnbaugh, Yuan Tian, Darren Dumlao, Renuka Nayak, Andrew Patterson, John C. Newman, Peter A. Crawford, Peter J. Turnbaugh

## Abstract

Diet can protect from autoimmune disease; however, whether diet acts via the host and/or microbiome remains unclear. Here, we use a ketogenic diet (KD) as a model to dissect these complex interactions. A KD rescued the experimental autoimmune encephalomyelitis (EAE) mouse model of multiple sclerosis in a microbiota-dependent fashion. Dietary supplementation with a single KD-dependent host metabolite (β-hydroxybutyrate, βHB) rescued EAE whereas transgenic mice unable to produce βHB in the intestine developed more severe disease. Transplantation of the βHB-shaped gut microbiota was protective. *Lactobacillus* sequence variants were associated with decreased T helper 17 (Th17) cell activation *in vitro*. Finally, we isolated a *L. murinus* strain that protected from EAE, which was phenocopied by the *Lactobacillus* metabolite indole lactic acid. Thus, diet alters the immunomodulatory potential of the gut microbiota by shifting host metabolism, emphasizing the utility of taking a more integrative approach to study diet-host-microbiome interactions.

## INTRODUCTION

Diet has broad impacts on autoimmune disease, including multiple sclerosis (MS), inflammatory bowel disease, and rheumatoid arthritis^1–5^, but the mechanisms responsible remain unclear. Consumption of a high-fat, low-carbohydrate ketogenic diet (KD) improves MS-related symptoms in humans^6–8^ and mice^9–11^. KDs are defined by a metabolic shift in host metabolism to lipid oxidation, resulting in elevated concentrations of the ketone bodies β-hydroxybutyrate (βHB) and acetoacetate (AcAc) in circulation^12^. AcAc and βHB have broad impacts on immune cells, acting to dampen inflammasome activation^13^, alter macrophage metabolism^14^, promote T cell function^15^, among other immunomodulatory effects^12^, providing a direct link between diet-induced shifts in host metabolites and the immune system.

However, KDs can also indirectly shape the immune system by altering the trillions of microorganisms that colonize the gastrointestinal (GI) tract (the gut microbiota) and/or their aggregate genomes and metabolic activities (the gut microbiome)^16–19^. KD-induced shifts in the gut microbiotas of humans and mice result in decreased intestinal immune activation in part due to the direct antimicrobial properties of βHB^16^. Yet the relevance of these diet-induced shifts in the gut microbiota for disease, especially diseases outside the gut that affect the brain or other organs, remained unknown.

Herein, we dissect the role of diet in shaping host-microbiome interactions relevant to disease. We chose to focus on the experimental autoimmune encephalomyelitis (EAE) mouse model for multiple reasons. The EAE model has been widely used in the autoimmunity field^20,21^, is microbiota-dependent^22^, and can be readily performed in gnotobiotic mice^23^. While human MS differs in many ways from EAE^21^, multiple MS-associated microorganisms exacerbate the EAE phenotype^22–24^ and transplantation of the gut microbiota from MS patients increases disease in the EAE model relative to healthy controls^23,25^. These results indicate that aspects of EAE reflect clinically relevant host-microbiome interactions, providing mechanistic insights essential to interpret the large-scale studies correlating aspects of the gut microbiome with MS in patients^26^.

We identify a pathway through which a microbial community shaped by intestinal production of βHB protects from EAE disease. By pairing recently described methods for generating stable in vitro communities^27^ with a Th17 skewing assay, we identify and isolate an immunomodulatory member of the *Lactobacillus murinis* species from a mouse fed the KD. This strain and its metabolite indole-3-lactate (ILA) were sufficient to decrease immune activation and ameliorate the EAE phenotype. These results provide new targets for therapeutic manipulation and a foundation to explore the mechanistic links between host and microbial metabolism in response to defined dietary perturbations.

## RESULTS

### The impact of diet on neurological phenotypes is microbiota-dependent

First, we sought to confirm that a ketogenic diet (KD) protects conventionally-raised (CONV-R) mice from EAE^9,10^. We fed 12-week-old female C57BL/6J CONV-R (*i.e.,* specific pathogen free, SPF) mice a high-fat diet (HFD, 75:15:10% fat:carbohydrate:protein) or a matched-ingredient KD (90.5:0:9.5% fat:carbohydrate:protein; diet details in **Table S1**; n=14 mice/group). Our diets were specifically formulated to control for the differences in dietary ingredients that have confounded previous comparisons of semi-purified KDs and the more complex chow diet^10,11,16,28–30^. As expected, the KD led to significantly higher levels of circulating βHB relative to HFD controls (**Figures S1A**).

Ten days after diet initiation we induced EAE by immunizing mice with the myelin oligodendrocyte glycoprotein (MOG) peptide. Disease was assessed by four distinct metrics commonly used in the autoimmunity field^22,24,31^: (*i*) changes in disease score over time; (*ii*) the overall rate of disease incidence; (*iii*) the maximum disease developed; and (*iv*) the distribution of mice across maximum disease scores. As EAE is known to present differently between experiments^20,31^, we included these same metrics in all experiments for the purpose of transparency regardless of baseline disease severity.

The KD significantly reduced disease severity. While HFD mice gradually increased in disease score over 28 days, the KD group had lower disease severity over time (**Figure 1A**). The incidence rate for the HFD group was 60% compared to 40% in the KD group (**Figure 1B**) with an average maximum disease score of diseased mice of 3.81±0.38 compared to 2.68±0.63 in the KD group (**Figures 1C,D**). These symptoms were consistent with flow cytometry-based analyses of immune cells in the brain and spleen. We assessed immune cell levels at 16 days post-immunization via flow cytometry. KD fed mice had significantly lower levels of CD4+ helper T cells co-producing IFNγ and IL-17a in the brain and spleen, which have been previously implicated in EAE^32^ (**Figures 1E-H**; gating strategy and total numbers in **Figure S1B-D**). Together with prior data from the intestine of healthy mice^16^, these results indicate that the impact of the KD on immune activation extends beyond the GI tract to alter systemic immunity and even the central nervous system (CNS).

**Figure 1.**
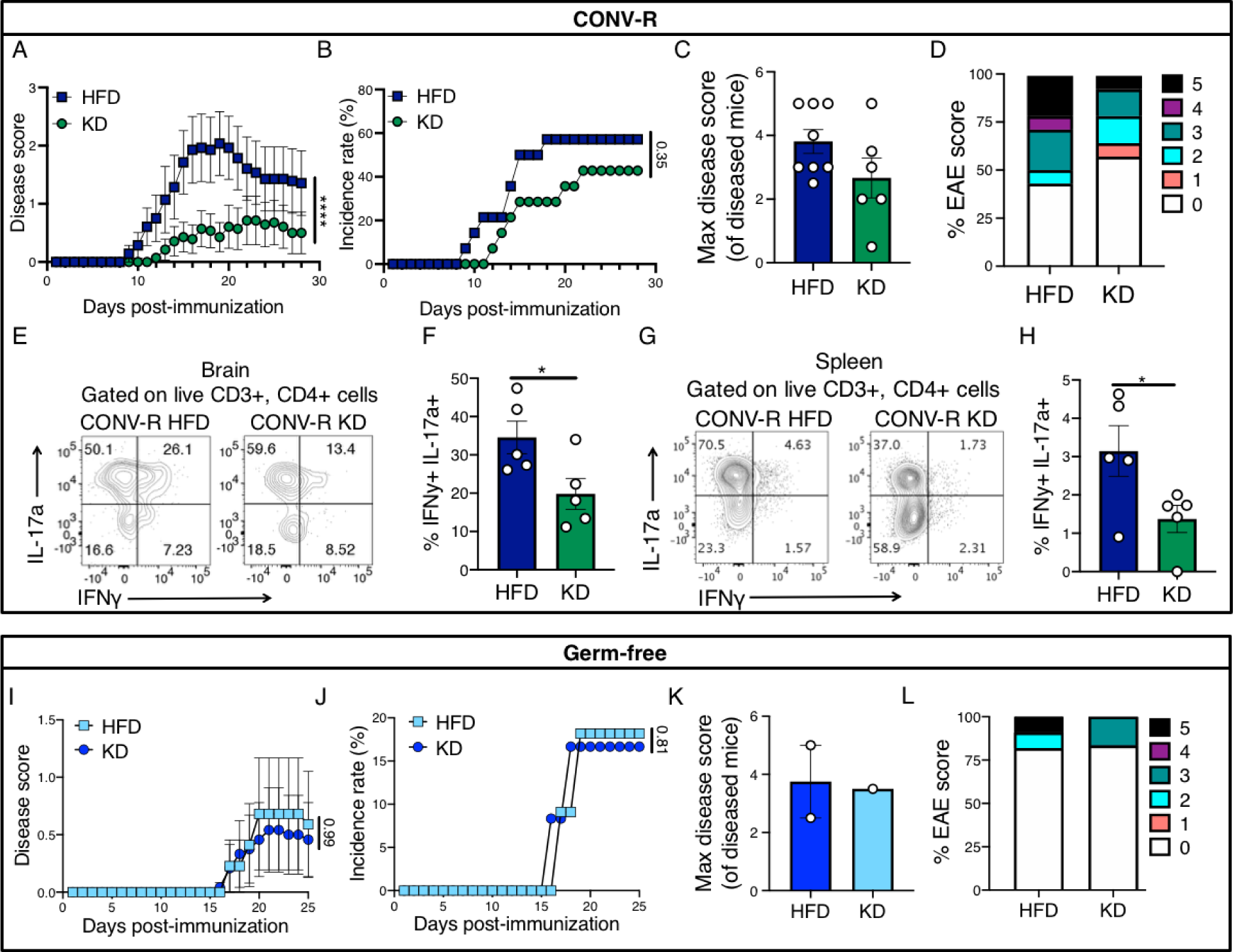
A ketogenic diet rescues a mouse model of multiple sclerosis in a microbiota-dependent manner. (A-D) Conventionally raised (CONV-R), specific pathogen free, female C57BL/6 Jackson mice were fed a high-fat diet (HFD) or a ketogenic diet (KD) (**Table S1**) for 10 days prior to induction of the experimental autoimmune encephalomyelitis (EAE) mouse model of multiple sclerosis (MS) (n=14/group; two independent experiments). (A) Disease scores (\*\*\*\**p*-value<0.0001; two-way ANOVA; mean±SEM) and (B) disease incidence (*p*-value listed; Log-rank Mantel-Cox test; percentage) were tracked over 28 days post-immunization. (C) Mean disease severity of mice that develop disease (mean±SEM; each point represents an individual mouse). (D) The proportion of mice with max disease score after 28 days. (E-H) At day 16 post-immunization mice fed a HFD or KD for 4 days prior to EAE induction (n=5/group) were sacrificed and brain and splenic lymphocytes were profiled for T helper subsets. (E) Representative flow cytometry from brain lymphocytes of IL-17a and IFNγ cells within the live CD3+ CD4+ population. (F) Percentage of IL-17a+ IFNγ+ cells within the live CD3+ CD4+ population in the brain. (\**p*-value<0.05, Welch’s t test; mean±SEM; each point represents an individual mouse) (G) Representative flow cytometry from splenocytes of IL-17a and IFNγ cells within the live CD3+ CD4+ population. (H) Percentage of IL-17a+ IFNγ+ cells within the live CD3+ CD4+ population in the spleen. (\**p*-value<0.05; Welch’s t test; mean±SEM; each point represents an individual mouse). (I-L) Germ-free (GF) female C57BL/6J mice were fed a HFD or KD for 7 or 16 days before EAE was induced and (I) disease scores (*p*-value listed; two-way ANOVA; mean±SEM) and (J) incidence rates (*p*-value listed; Log-rank Mantel-Cox test; percentage) were tracked for 25 days post-immunization (n=11 HFD; n=12 KD; two independent experiments). (K) Mean maximum disease severity of mice that developed disease (mean±SEM; each point represents a diseased mouse). (L) The proportion of mice with max disease score after 25 days.

Next, we sought to test if the protective effect of a KD during EAE is dependent on the microbiota. We repeated the same dietary intervention using 9-12-week-old C57BL/6J female germ-free (GF) mice (n=11-12 mice/group; 4 cages/isolator; 2 independent experiments). Mice were fed a HFD or KD for 7 days prior to EAE induction, as described above, and monitored for 25-32 days. Serum βHB was significantly elevated in GF KD fed mice relative to HFD controls (**Figure S1C**). However, in contrast to CONV-R mice (**Figures 1A-D**), we did not detect any significant differences between GF mice fed HFD and KD across any of the 4 disease metrics (**Figures 1I-L**). Consistent with prior work^22^, GF mice on the HFD had significantly decreased disease phenotypes relative to HFD fed CONV-R mice (**Figure S1D**; 0% incidence for GF mice at day 16 vs. 80% for CONV-R). Thus, the microbiota alters both the initiation of disease and the protective effect of a KD.

### Intestinal βHB is necessary and sufficient to protect from disease

The dietary shift to a KD has broad implications for host and microbial metabolism that could feasibly alter the immune system^33^. We opted to focus on the ketone body β-hydroxybutyrate (βHB) (**Figure 2A**) given its dependence on the KD and its prior links to both immune cells^34–36^ and the microbiota^16^. We supplemented the HFD with a βHB-containing ketone ester (100 g/kg hexanoyl hexyl β-hydroxybutyrate, C6×2-βHB, see HFD-KE in **Table S1**) which leads to increased gut and circulating βHB^16^. We fed 12-week-old CONV-R male and female C57BL/6J mice a HFD, HFD-KE or KD for 3 days prior to EAE disease induction (n=9-10 mice/group). Disease was tracked for 21 days prior to euthanasia.

**Figure 2.**
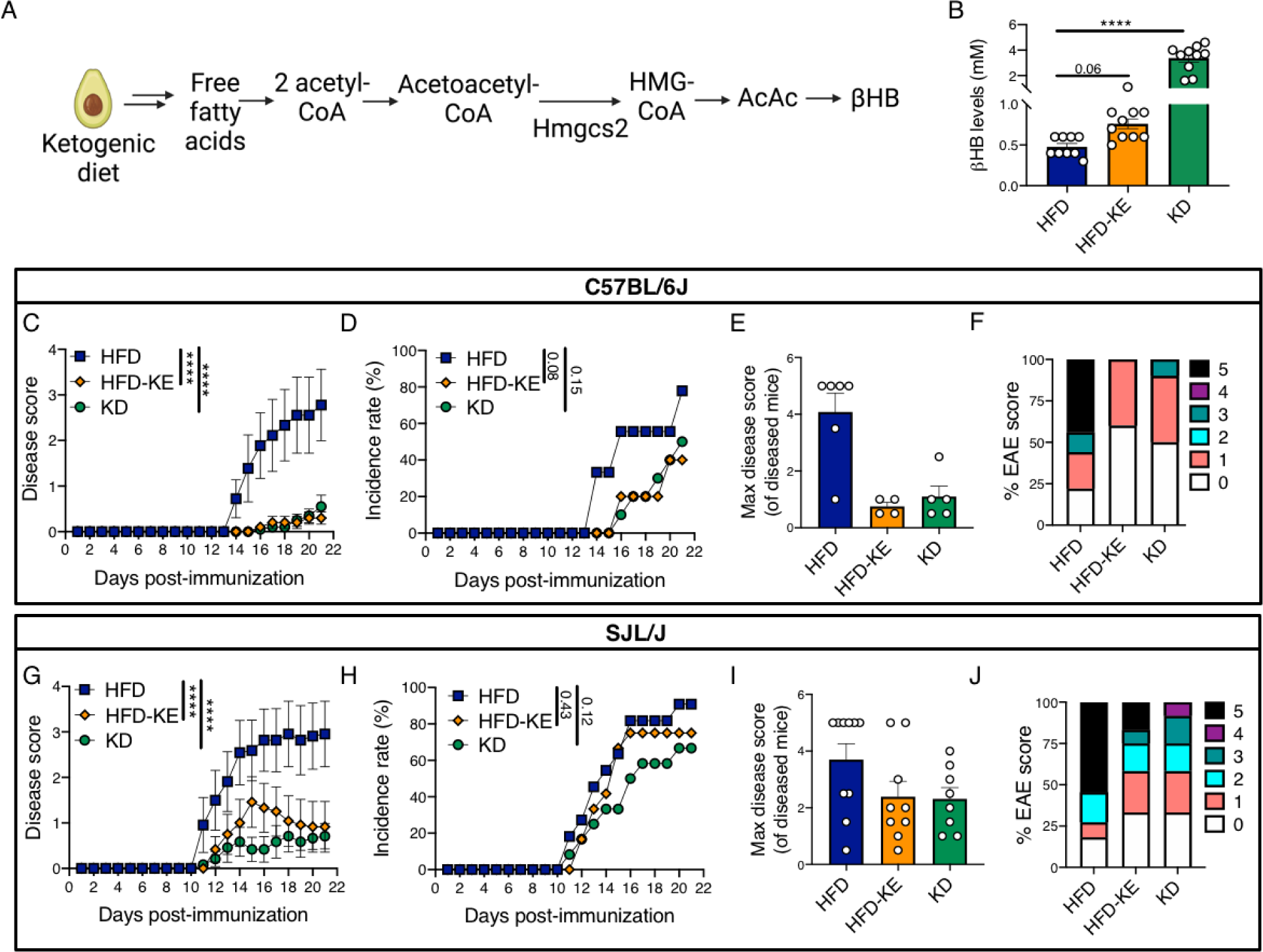
Intestinal βHB is sufficient for protection from neurological disease. (A) Pathway for β-hydroxybutyrate (βHB) production where 3-Hydroxy-3-Methylglutaryl-CoA Synthase 2 (Hmgcs2) is the rate-limiting enzyme. (AcAc: acetoacetate) (B-F) Conventionally raised (CONV-R), specific pathogen free, male and female C57BL/6J jackson mice were fed a high-fat diet (HFD), a HFD supplemented with a βHB ketone ester (HFD-KE), or a ketogenic diet (KD) for 3 days prior to EAE disease induction (**Table S1**). (n=9 HFD; n=10 HFD-KE and KD) (B) Circulating βHB levels 7 days post-immunization. (C) Disease scores (\*\*\*\**p*-value<0.0001; listed; two-way ANOVA; mean±SEM) and (D) disease incidence (*p*-value listed; Log-rank Mantel-Cox test; percentage) was tracked over 21 days post-immunization. (E) Mean maximum disease severity of mice that developed disease (mean±SEM; each point represents a diseased mouse). (F) The proportion of mice with max disease score after 21 days. (G-J) Conventionally raised (CONV-R), specific pathogen free, female SJL Jackson mice were fed a HFD, HFD-KE, or a KD for 5 days prior to EAE disease induction (n=11 HFD; n=12 HFD-KE and KD). (G) Disease scores (\*\*\*\**p*-value<0.0001; listed; two-way ANOVA; mean±SEM) and (H) disease incidence (*p*-value listed; Log-rank Mantel-Cox test; percentage) was tracked over 21 days post-immunization. (I) Mean maximum disease severity of mice that developed disease (mean±SEM; each point represents a diseased mouse). (J) The proportion of mice with max disease score after 21 days.

Remarkably, the pharmacological administration of the βHB ketone ester (βHB-KE) was sufficient to protect mice from EAE. Consistent with our previous findings^16,37^, circulating βHB levels were increased with βHB-KE supplementation and KD (**Figure 2B**). In line with our previous results (**Figure 1A**), the HFD group had a significantly increased disease score over time relative to the KD group (**Figure 2C**). The HFD-KE group was indistinguishable from KD by either disease score (**Figure 2C**) or incidence rate (**Figure 2D**). The average maximum disease scores of diseased mice were markedly lower in HFD-KE (0.75±0.14) and KD (1.10±0.37) relative to HFD (4.08±0.66, **Figure 2E**) with 100% of the HFD-KE mice remaining at a score of ≤2 (**Figure 2F**). We additionally investigated if the KD and KE supplementation were protective in a relapsing-remitting course of EAE using the SJL mouse model with proteolipid protein (PLP)^38,39^. A KD and HFD-KE were both protective in the SJL mouse of EAE, where both KD and HFD-KE had decreased disease scores over time compared to HFD fed mice with similar incidence rates to HFD (**Figure 2G-H**). The maximum disease scores of diseased mice were lower in KD and HFD-KE (**Figure 2I-J**). In conclusion, βHB-KE supplementation is sufficient to improve EAE disease severity.

Although we were orally delivering βHB through the diet, it could potentially exert an impact on EAE directly in the central nervous system (CNS) or in other tissues following absorption from the GI tract. To control for this effect, we generated and validated a transgenic mouse model to selectively abolish the intestinal production of βHB (**Figure S2**). 3-Hydroxy-3-Methylglutaryl-CoA Synthase 2 (HMGCS2) is the rate-limiting enzyme for βHB production and is highly expressed in intestinal epithelial cells (IECs) within the large intestine^40^. We generated *Hmgcs2^WT^*(*Hmgcs2^fl/fl^ VillinER/Cre^-/-^*) and *Hmgcs2^ΔIEC^* (*Hmgcs2^fl/fl^ VillinER/Cre^+/-^*) mice (**Figure S2A**). Following tamoxifen administration, HMGCS2 was undetectable in colonic IECs by Western blot with no change in the liver (**Figure S2B-C**). βHB was nearly undetectable in colonic IECs from *Hmgcs2^ΔIEC^*mice (**Figure S2D**). In contrast, serum βHB was unaffected (**Figure S2E**), suggesting that the direct impact of intestinal ketogenesis is restricted to the GI tract.

Consistent with our KE experiment, we found that HMGCS2 in IECs is required for KD-mediated protection during EAE. We treated 7-9-week-old CONV-R male *Hmgcs2^ΔIEC^* and *Hmgcs2^WT^* mice with tamoxifen daily for 5 days, allowed for a 5 day washout, then put the mice on a KD for 3 days followed by EAE induction (**Figure S3A**; n=10-11 mice/group; 2 independent experiments). *Hmgcs2^ΔIEC^* mice had significantly increased disease scores over time relative to their *Hmgcs2^WT^* counterparts (**Figure 3A**). Disease incidence (**Figure 3B**) and maximum disease score (**Figure 3C**) were lower in *Hmgcs2^WT^* mice. The range of disease scores was also markedly different between *Hmgcs2^WT^* (score=0-2) and *Hmgcs2^ΔIEC^*(score 0-4; **Figure 3D**). Importantly, circulating βHB levels were not altered between the *Hmgcs2^WT^* and *Hmgcs2^ΔIEC^* mice (**Figure 3E**). These phenotypic differences were consistent with the significantly higher IL-17a+ IFNγ+ Th17 cells within the brains of *Hmgcs2^ΔIEC^* mice relative to controls (**Figures 3F-G** and **Figure S3B**; n=10-11 mice/group; 2 independent experiments). IL-17a mean fluorescence intensity was also significantly higher in *Hmgcs2^ΔIEC^*mice (**Figure 3H**), indicative of increased Th17 cell activation.

**Figure 3.**
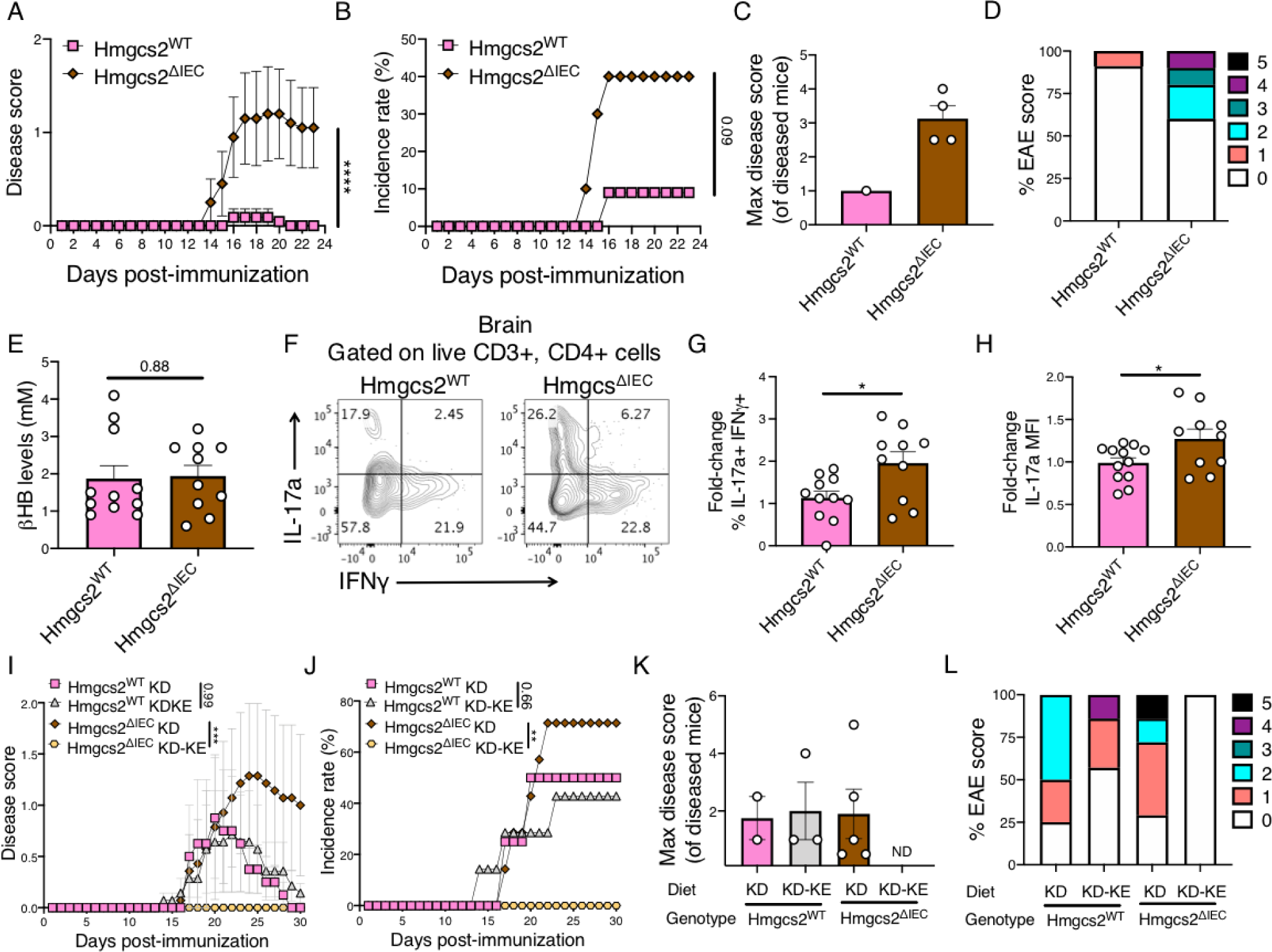
Intestinal βHB is necessary for protection from neurological disease. (A-H) Male *Hmgcs2^WT^* (*Hmgcs2^fl/fl^ VillinER/Cre^-/-^*) and mice with *Hmgcs2* specifically deleted from intestinal epithelial cells (*Hmgcs2^ΔIEC^ - Hmgcs2^fl/fl^ VillinER/Cre^+/-^*) were injected with tamoxifen daily for 5 days in induce *Hmgcs2* deletion. Mice then had a washout period of 5 days before being given with a KD for 3 days before EAE was induced (n=11 *Hmgcs2^WT^*; n=10 *Hmgcs2^ΔIEC^*; 2 independent experiments). (A) Disease scores (\*\*\*\**p*-value<0.0001; two-way ANOVA; mean±SEM) and (B) disease incidence (*p*-value listed; Log-rank Mantel-Cox test; percentage) was tracked for 23 days. (C) Mean maximum disease severity of mice that developed disease (mean±SEM; each point represents a diseased mouse). (D) The proportion of mice with max disease score after 23 days. (E) Circulating βHB levels 7 days post-immunization. (F-H) After 23 days mice were sacrificed and levels of Th17 cells in the brain were quantified with flow cytometry. (F) Representative flow cytometry from brain lymphocytes of Th17 cells (IL-17a+ IFNγ+ cells within the live CD3+ CD4+ population). (G) Fold-change relative to the *Hmgcs2^WT^* group in the IL-17a+ IFNγ+ cells within the live CD3+ CD4+ population and (H) IL-17a mean fluorescence intensity (MFI) within the live TCRβ+ population (\**p*-value<0.05; Welch’s t test; mean ± SEM; each point represents an individual mouse). (I-L) Male *Hmgcs2^WT^* (*Hmgcs2^fl/fl^ VillinER/Cre^-/-^*) or *Hmgcs2^ΔIEC^* (*Hmgcs2^fl/fl^ VillinER/Cre^+/-^*) were injected with tamoxifen daily for 5 days then mice had a washout period of 5 days before put on a KD or a KD supplemented with a βHB ketone ester (KD-KE) (**Table S1**) for 5 days before EAE was induced. (n=4 *Hmgcs2^WT^* KD, n=7 *Hmgcs2^WT^* KD-KE, *Hmgcs2^ΔIEC^* KD and *Hmgcs2^ΔIEC^*KD-KE) (I) Disease scores (\*\*\**p*-value<0.001; listed; two-way ANOVA; mean±SEM) and (J) disease incidence rates (\*\**p*-value<0.01 or listed; Log-rank Mantel-Cox test; percentage) were tracked for 30 days. (K) Mean maximum disease severity of mice that developed disease (mean±SEM; each point represents a diseased mouse; ND = not detected). (L) The proportion of mice with max disease score after 30 days.

Finally, we used the pharmacological administration of the KE to rescue disease caused by the genetic deletion of *Hmgcs2* in IECs. We fed 14-17-week-old CONV-R male *Hmgcs2^ΔIEC^*and *Hmgcs2^WT^* mice a KD or the same KD supplemented with the KE (100 g/kg, **Table S1;** n=4-7 mice/group). After 3 days on diet we induced EAE and tracked disease scores for 30 days. Supplementation of the KE significantly increased circulating βHB levels in both *Hmgcs2^ΔIEC^* and *Hmgcs2^WT^* mice compared to unsupplemented KD (**Figure S3C**). As expected, *Hmgcs2^WT^*mice were unaffected by the addition of the KE (**Figures 3I-L**). In contrast, the KE significantly decreased disease scores over time in *Hmgcs2^ΔIEC^* mice (**Figure 3I**) and incidence rate (**Figure 3J**). Maximum disease scores of diseased mice were also lower in the KE supplemented *Hmgcs2^ΔIEC^*mice relative to KD controls (**Figure 3K**), with 100% of the KD-KE fed *Hmgcs2^ΔIEC^* mice having no development of clinical disease by day 30 (**Figure 3L**). *En masse*, these experiments all support the key role of the intestinal production of βHB for the protective effect of a KD during EAE.

### The antibiotic ampicillin phenocopies the effects of the ketone ester

Given the potential for βHB to alter numerous host^12,41^ and microbial^16^ cells within the GI tract, we sought to test if the effects of the KE are microbiota-dependent. Prior work had demonstrated that the broad-spectrum beta-lactam antibiotic ampicillin rescues EAE in mice fed a standard chow diet^24^; however, the effects on a HFD or KD were unknown. We fed 9-week-old CONV-R female C57BL/6J mice a HFD or a matched HFD-KE for 3 days prior to administration of ampicillin in their drinking water (n=6-7 mice/group). Following 7 days of ampicillin treatment we induced EAE and tracked disease for 25 days (**Figure S4A**). Replicating our prior experiment (**Figures 2C-F**), HFD-KE mice had significantly lower disease scores than the HFD group in the absence of ampicillin (**Figure S4B**). Consistent with the prior literature^24^, ampicillin significantly decreased EAE disease scores over time in HFD fed mice (**Figure S4B**), with a trend towards lower disease incidence (**Figure S4C,** *p*=0.06, Log-rank Mantel-Cox test). In contrast, HFD-KE fed mice treated with or without ampicillin were indistinguishable from each other (**Figures S4B-C**). The maximum scores in diseased mice were comparable between groups with a slight reduction in HFD mice treated with ampicillin (**Figures S4D-E**). Combined, these results indicate that the protective effects of βHB are microbiota-dependent, prompting us to revisit the impact of diet and KE supplementation on gut microbial community structure.

### Ketogenesis alters the gut microbiotas of diseased mice

We hypothesized that intestinal βHB could protect from EAE in part due to altering the immunomodulatory potential of the gut microbiota. However, our prior studies were performed in healthy mice^16^, so it remained unclear if the KD and KE would have significant impacts on the gut microbiota in diseased mice. We used 16S rRNA gene sequencing (16S-seq) to profile the gut microbiotas of mice at 7 days post-immunization from the same HFD, HFD-KE, and KD fed EAE mice in **Figure 2B-F**. 16S-seq revealed significant shifts in the gut microbiota between the three diets during EAE (**Figure S5**).

Shannon diversity was higher in the HFD-KE and KD groups compared to the HFD (**Figure S5A**). Principal coordinate analysis (PCA) euclidean distances with PERMANOVA testing revealed a significant difference between HFD-KE and HFD as well as KD and HFD (**Figure S5B**). The KE and the KD led to largely consistent changes in the abundance of <u>a</u>mplicon <u>s</u>equence <u>v</u>ariants (ASVs) relative to the HFD, with some notable exceptions (**Figure S5C,D**). Thus, diet and exogenous delivery of βHB still have a widespread impact on the composition of the gut microbiota in diseased mice.

### The βHB-associated gut microbiota protects from disease

Having shown that βHB has a broad impact on the gut microbiota, we next tested the impact of fecal microbiota transplantation (FMT) on EAE phenotypes. We fed 6-8-week-old female CONV-R C57BL/6J mice a HFD or HFD-KE for 3 days prior to receiving a commonly used antibiotic cocktail (AVNM; <u>a</u>mpicillin, <u>v</u>ancomycin, <u>n</u>eomycin, and <u>m</u>etronidazole) for 1 week to deplete the baseline microbiota^42,43^. Fecal microbiota transplantation (FMT) was performed daily by oral gavage for 7 days prior to and after EAE induction from 6-8-week-old CONV-R female donor mice fed a HFD or HFD-KE for 1-3 weeks (2 donors, 4 recipients/donor, 2 cages/group; **Figure 4A**).

**Figure 4.**
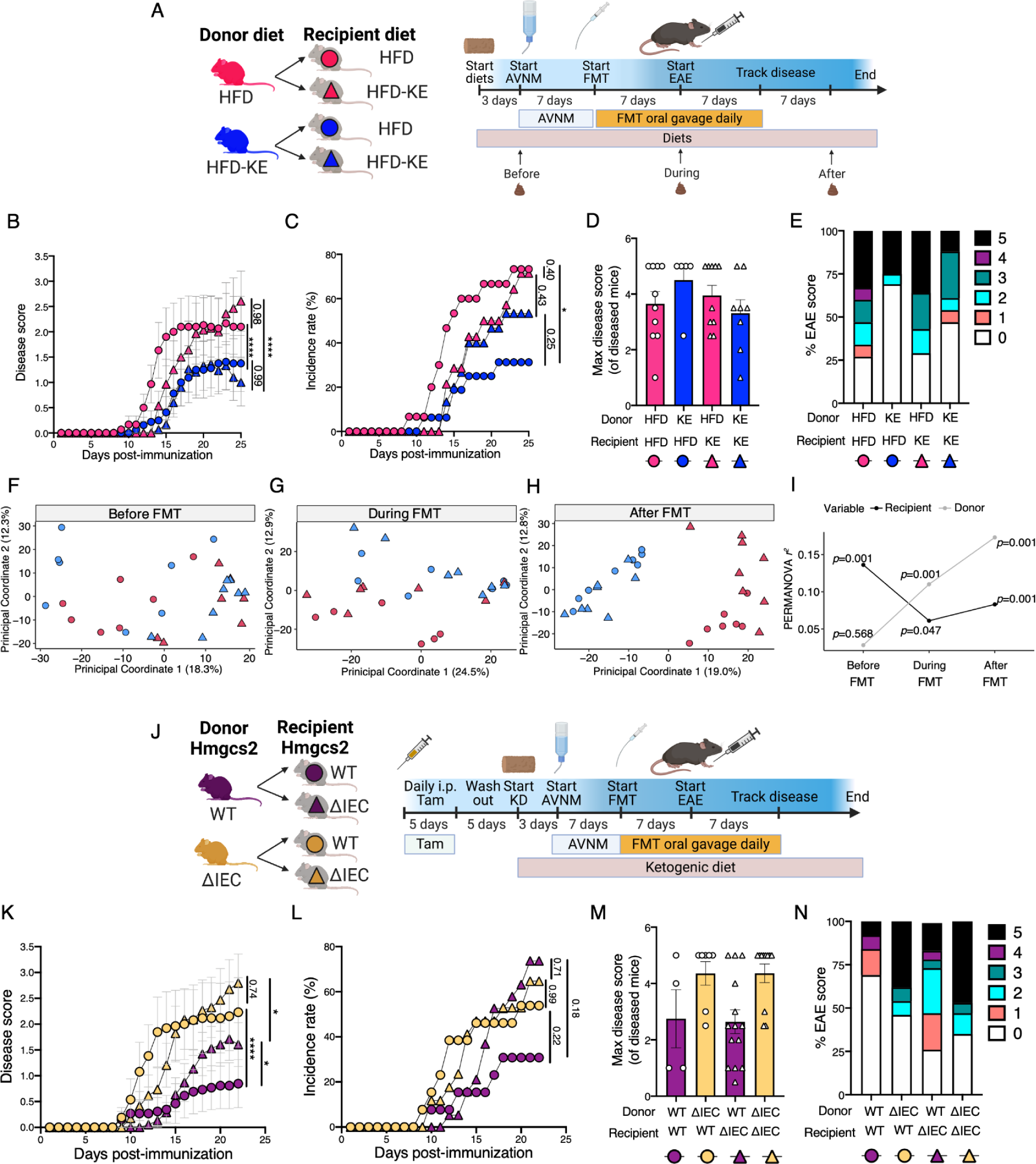
The βHB-altered gut microbiota is sufficient to protect from disease. (A-I) Fecal microbiota transplantation experiments between high-fat diet (HFD) or a HFD supplemented with a βHB ketone ester (HFD-KE) fed mice. HFD or HFD-KE donor fecal microbiota were transplanted into AVNM treated HFD or HFD-KE fed recipients then EAE was induced (n=15 HFD microbiota to HFD recipient - pink circles; n=14 HFD microbiota to HFD-KE recipient - pink triangles; n=16 HFD-KE microbiota to HFD recipient - blue circles; n=15 HFD-KE microbiota to HFD-KE recipient - blue triangles; two independent experiments). Timepoints for fecal sample collection for 16S-seq are indicated on the timeline. (A) Experimental design and timeline for diet FMT experiments. (B) Disease scores (\*\*\*\**p*-value<0.0001; two-way ANOVA; mean±SEM) and (C) disease incidence rates (\**p*-value<0.05 or listed; Log-rank Mantel-Cox test; percentage) were tracked over 25 days post-immunization. (D) Mean disease severity of mice that developed disease (mean±SEM; each point represents a diseased mouse). (E) The proportion of mice with max disease score after 25 days. (F-H) Principal coordinate analysis (PCA) of Euclidean distances (F) before, (G) during, and (H) after FMTs. (I) PERMANOVA testing of recipient (black) and donor (gray) diet before, during, or after FMTs. (J-N) *Hmgcs2^WT^* or *Hmgcs2^ΔIEC^* fecal microbiota was transplanted into KD-fed AVNM treated *Hmgcs2^WT^* and *Hmgcs2^ΔIEC^* recipient mice then EAE was induced. (n=13 *Hmgcs2^W^*^T^ microbiota to *Hmgcs2^WT^* recipient - purple circles, n=13 *Hmgcs2^ΔIEC^*microbiota to *Hmgcs2^WT^* recipient - purple triangles, n=19 *Hmgcs2^WT^* microbiota to *Hmgcs2^ΔIEC^* recipient - yellow circles, n=17 *Hmgcs2^ΔIEC^* microbiota to *Hmgcs2^ΔIEC^* recipient - yellow triangles). (J) Experimental design and timeline for *Hmgcs2* FMT experiments. (K) Disease scores (\*\*\*\**p*-value<0.0001; listed; two-way ANOVA; mean±SEM) and (L) disease incidence rates (*p*-value listed; Log-rank Mantel-Cox test; percentage) were tracked over 22 days post-immunization. (M) Mean disease severity of mice that developed disease (mean±SEM; each point represents a diseased mouse). (N) The proportion of mice with max disease score after 22 days.

FMT from HFD-KE fed donors significantly decreased EAE phenotypes relative to a HFD donor irrespective of the diet of the recipient mice. Disease scores over time were significantly lower in both recipient groups given the HFD-KE FMT relative to HFD FMT controls fed the same diet (**Figure 4B**). Incidence rate was also significantly lower in HFD fed recipients given an FMT from HFD-KE relative to HFD (**Figure 4C**). However, maximum disease scores (**Figure 4D**) and the distribution of scores (**Figure 4E**) were not markedly different between groups, suggesting that differences are driven by time to onset of disease and time to the development of severe disease. 16S-seq analysis of the recipient mice revealed that the donor gut microbiota had a significant impact on the gut microbiota that overwhelmed the baseline differences by recipient diet (**Figure 4F-I**). This major effect of the donor gut microbiota was likely driven by our use of repeated oral gavage for two weeks. The protective effect of the KE was less dramatic than in our earlier experiments (**Figure 2**), potentially due to the antibiotic pre-treatment and/or the normal variation in disease severity between experiments^20,31^.

Next, we performed a similar FMT experiment using our transgenic mice deficient in intestinal βHB production. Donor *Hmgcs2^WT^* and *Hmgcs2^ΔIEC^* mice (2 donors/group) were fed a KD for 3 days prior to FMT. Recipient mice of both genotypes were given AVNM via drinking water for 7 days prior to daily gavage for 1 week prior to and following EAE induction (**Figure 4J**). Disease was tracked for 22 days. Consistent with our KE FMT experiment (**Figures 4B-E**), we found that FMT from an *Hmgcs2^ΔIEC^* donor led to significantly higher disease scores irrespective of the genotype of the recipient mice (**Figure 4K**). Disease incidence was similar in most groups with the exception of a trending lower incidence in the *Hmgcs2^WT^* FMT to *Hmgcs2^WT^*recipient (**Figure 4L**). Maximum disease scores of mice that developed disease were lower in mice that received *Hmgcs2^WT^* FMT compared to *Hmgcs2^ΔIEC^*FMT with 38% and 47% of the *Hmgcs2^ΔIEC^* recipients developed maximum disease (**Figure 4M-N**).

Unlike the diet FMT experiment, some effects of host genotype persisted despite having distinct FMT donors. *Hmgcs2^WT^* FMT into *Hmgcs2^WT^* recipients led to significantly lower disease scores over time compared to *Hmgcs2^WT^* FMT into *Hmgcs2^ΔIEC^* mice (**Figure 4K**). However, irrespective of the recipient genotype *Hmgcs2^ΔIEC^* FMT resulted in worsened disease over time (**Figure 4K**). The combined results across both FMT paradigms suggest that intestinal βHB alters the gut microbiota in a manner that ameliorates the EAE model.

### Identification of immunomodulatory bacteria linked to diet

Our prior work demonstrated a reproducible decrease in bifidobacterial abundance in both humans and mice fed a ketogenic diet, resulting in decreased intestinal Th17 activation^16^. However, we only detected a single ASV within the *Bifidobacterium* genus within the gut microbiota of the mice in our initial experiment (**Figure S5**), which was not significantly altered in response to diet (0.4%±0.19% HFD vs. 0.3%±0.11% KD, relative abundance, *p*=0.75, Welch’s t test). Furthermore, a pilot experiment in which we orally gavaged *Bifidobacterium adolescentis* strain BD1 to CONV-R mice fed a ketogenic diet did not significantly impact the EAE phenotype (*data not shown*), prompting us to design an unbiased approach to screen for bacteria of interest that could mediate the protective effect of a KD.

Given that previous studies have implicated a range of microorganisms capable of impacting EAE^44^, we aimed to take a reductionist approach to narrow in on members of the microbiota contributing to the protective effect of a KD and KE supplementation. We utilized a method for generating <u>s</u>table *in vitro* <u>c</u>ommunities (SICs) that was initially established using the human gut microbiota^45^. Stool samples from a representative mouse fed the HFD, HFD-KE, or KD collected 7 days post-immunization were passaged 3 times in rich bacterial media (n=4-5 SICs/donor; see *Methods*; **Figure S6A**). 16S-seq was used to assess the microbial communities remaining in each SIC. Multiple differences between donor samples persisted despite 3 rounds of passaging *in vitro.* The KD SICs had significantly lower Shannon diversity than HFD SICs (**Figure S6B, Table S2**). Despite a marked divergence from the donor samples (**Figure S6C**), the SICs from the KD and KE donors were both significantly different from the HFD SICs (**Figure S6C,D**). Overall richness was quite low after passaging, with just 16 unique ASVs detected in the SICs at the endpoint (4-10 ASVs/SIC; **Figure 4E**) compared to 41, 109, and 128 ASVs in the donor HFD, HFD-KE, and KD samples respectively. Three ASVs were both abundant and common across all 13 SICs, including *Enterococcus* (ASV4), *Bacteroides* (ASV2), and *Parasutterella* (ASV9). The remaining ASVs were variable between samples and diet groups.

Th17 cells play an important role in the pathogenesis of both human MS and the EAE model^46–49^. Thus, we used our previously established Th17 skewing assay^50,51^ to assess the aggregate immunomodulatory potential of each SIC in the absence of other immune or non-immune cell types. Briefly, splenic CD4+ T cells were isolated and skewed to Th17 fate in the presence of cell free supernatants from our SICs. After 4 days, cells were then restimulated with PMA and ionomycin overnight before supernatants were collected for IL-17a quantification via ELISA. All of the HFD-derived SICs induced IL-17a whereas none of the KD-derived SICs had an effect in this assay (**Figure 5A, top**); the HFD-KE SICs were mixed, with 4/5 inducing IL-17a. Three ASVs were significantly associated with IL-17a levels (*p*-value<0.05, Spearman rank correlation; **Figure 5A, right**), including two negatively associated ASVs annotated as *Lactobacillus* (ASV3 and ASV24; **Figures 5B-C**) and the positively associated *Parasutterella* (ASV9; **Figure S6F**). Note, *Lactobacillus* was recently renamed *Ligilactobacillus*; however, we opted to refer to it as *Lactobacillus* for consistency with previous studies^52–54^.

**Figure 5.**
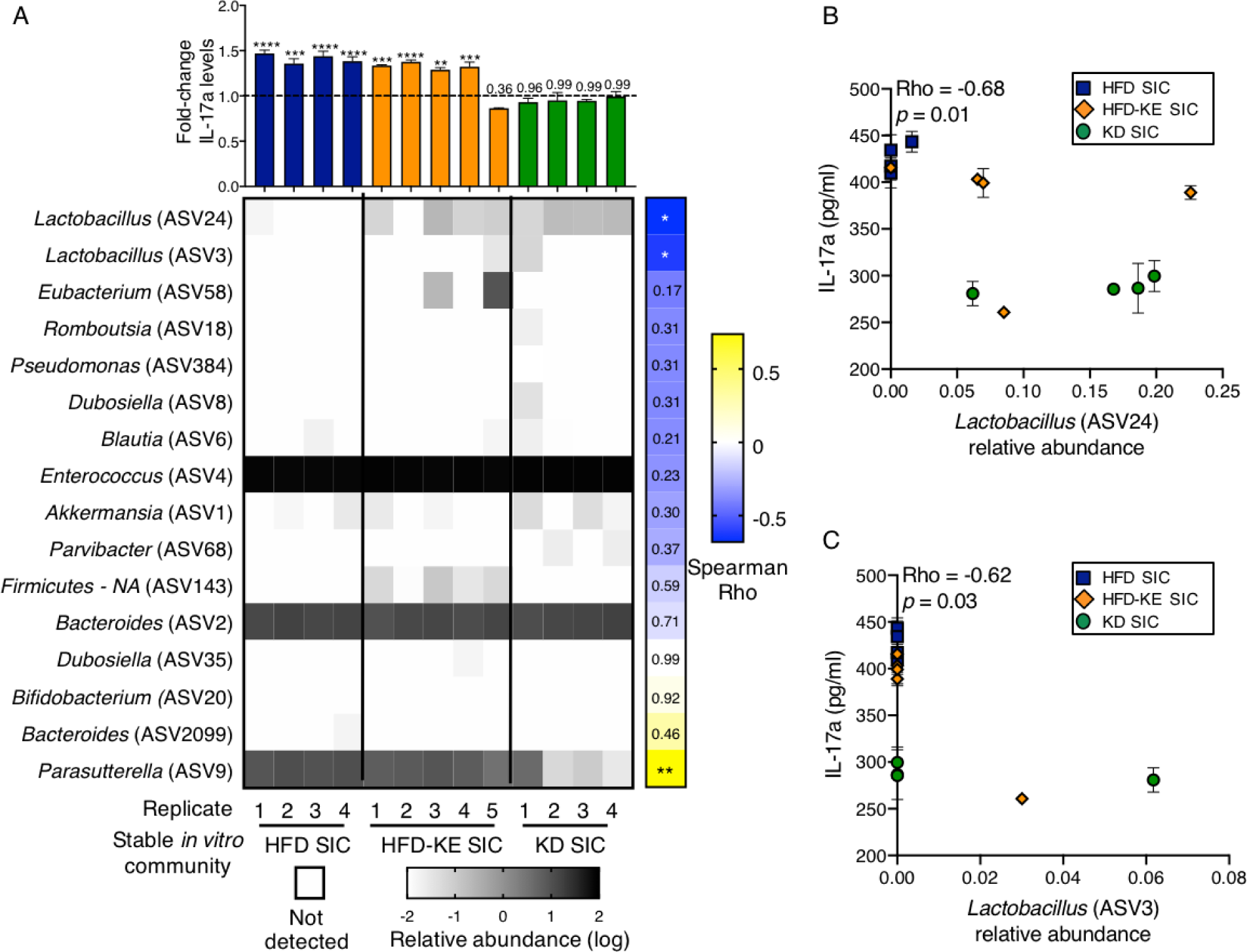
*Lactobacillus* sequence variants are associated with decreased Th17 activation. (A-C) Stool stable *in vitro* communities (SICs) were derived from high-fat diet (HFD) HFD supplemented with a βHB ketone ester (HFD-KE) or a ketogenic diet (KD) mouse stool 7 days post EAE induction (1 donor/group, 4-5 SIC replicates). Cell free supernatants from these SICs were tested for their capacity to alter Th17 skewing and community structure was assessed with 16S rRNA gene sequencing. (A) **Top -** IL-17a levels were measured via ELISA post re-stimulation of Th17 skewed splenic T cells treated with cell free supernatant from HFD, HFD-KE, and KD EAE derived SICs where cells were treated for 4 days. (n=3 for Th17 skewing biological replicates; \*\**p-*value<0.01; \*\*\**p-*value<0.001; \*\*\*\**p-*value<0.0001; listed one-way ANOVA with Dunnett’s multiple comparisons test compared to BHI media control; Fold-change is relative to the BHI media control which is represented by the dotted line; mean±SEM). **Bottom -** Heat map of the relative abundance (log) of amplicon sequence variants (ASVs) in the SICs with assigned genus (SILVA v138 database). White denotes the absence of an ASV. **Right -** Spearman correlations Rho values of ASV relative abundance and IL-17a levels (\**p-*value<0.05; \*\**p-*value<0.01; or listed; Spearman correlation). (B-C) Significantly negatively correlated ASVs with IL-17a levels, (B) ASV24 and (C) ASV3 from the Spearman correlation of ASV relative abundance and IL-17a levels are plotted in individual scatter plots (\**p-*value<0.05; \*\**p-*value<0.01; Spearman correlation; mean±SEM plotted for IL-17a levels).

### A *Lactobacillus* isolate and metabolite protect from disease

We sought to isolate a representative *Lactobacillus* strain from our SICs to enable more controlled experiments. We plated KD-SIC replicate 4 (**Figure 5A**) on *Lactobacillus*-selective media (MRS) prior to 2 days of anaerobic incubation. Identification by matrix assisted laser desorption/ionization (MALDI) mass spectrometry revealed 2 colonies that matched the *Lactobacillus murinus/animalis* fingerprint; a representative isolate was validated by full-length 16S-seq (**Table S3**). Multiple lines of evidence suggested that this isolate matched ASV24 from the SICs: (*i*) ASV24 was the only ASV assigned to *Lactobacillus* in the source SIC (KD SIC replicate 4) and (*ii*) the 16S rRNA gene sequence was 100% identical with 100% coverage (NCBI BLASTN) to the representative sequence from ASV24 (**Table S3**). We designated this isolate *Lactobacillus murinus* KD6.

Lactobacilli, including *L. murinus,* convert tryptophan (Trp) to indole-3-lactate (ILA), which inhibits Th17 cell production of IL-17a^53,55^. To assess if *L. murinus* KD6 had genes involved in ILA production, we generated a finished genome using a hybrid of short- and long-read sequencing (see *Methods*). Trp is deaminated via aromatic amino acid aminotransferases (ArATs) to form indole-3-pyruvate (IPYA) which is then dehydroxylated by phenyllactate dehydrogenases (FldH) or D-lactate dehydrogenases (D-LDH) to make ILA^52,54–56^ (**Figure 6A**). We identified 4 genes homologous to aminotransferases in the *L. murinus* KD6 genome. This includes a putative aspartate aminotransferase (AspAT) with significant homology to an annotated AspAT from known ILA producing species, including *Clostridium sporogenes, L. reuteri*, and *L. murinus* (**Figure 6B, Tables S4 and S5**). We also detected 4 putative *LDH* genes in the *L. murinus* KD6 genome, including a candidate D-lactate dehydrogenase (*D-LDH*) with significant homology to genes found in ILA producers (**Figure 6B, Tables S4 and S5**). We then used a targeted liquid chromatography mass spectrometry (LCMS) assay to confirm the production of ILA from *L. murinus* KD6 and the positive control strain *L. reuteri* ATCC23272^52^ (**Figure 6C**). As expected^53,55^, ILA inhibited the production of IL-17a by Th17 cells in our skewing assay (**Figure 6D**).

**Figure 6.**
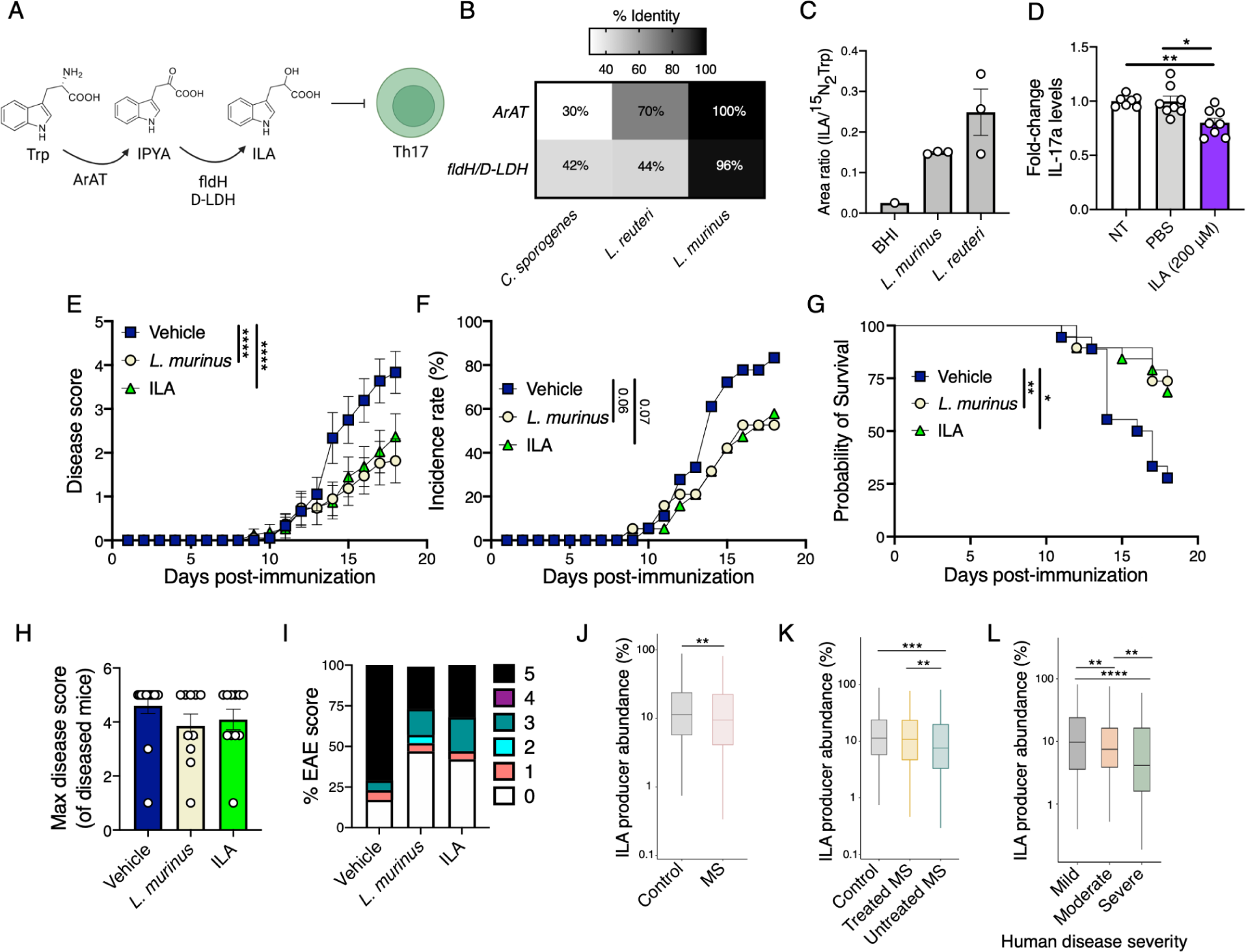
A *Lactobacillus murinus* isolate and metabolite decrease disease severity on a HFD. (A) Tryptophan (Trp) metabolism pathway where Trp is metabolized to indole-3-pyruvate (IPYA) via aromatic aminotransferase then IPYA is metabolized to ILA via indolelactate dehydrogenase (fldH) or lactate dehydrogenase (LDH). ILA has been shown to inhibit Th17 cells^45^. (B) Top hits in the *L. murinus* KD6 genome for ArAT and D-LDH with percent identity (listed) and compared to *Clostridium sporogenes, L. reuteri,* and *L. murinus* via BLASTP (*E*-value<1e^-51^). See also Tables S4 and S5. (C) ILA was quantified via high resolution liquid chromatography mass spectrometry from *L. murinus* KD6 or *L. reuteri* strain ATCC23272 cultured in BHI^CHV^. The area ratio of ILA to labeled Trp (^15^N_2_Trp) is displayed (n=3 biological replicates *L. murinus* KD6 and *L. reuteri;* n=1 for BHI control; mean±SEM). (D) IL-17a levels in Th17 skewed cells that were incubated for 4 days with no treatment (NT), a vehicle control (PBS), or 200 µM indole-3-lactate (ILA) post re-stimulation (n=7-8; each point represents a Th17 skewing biological replicate; \**p*-value<0.05 or listed; one-way ANOVA Kruskal-Wallis with Dunn’s test; mean±SEM; values are relative to the NT condition; two independent experiments). (E-I) Conventionally raised, specific pathogen free female Jackson mice were fed a high-fat diet (HFD) for 3 days prior to the start of every other day oral gavage with either a PBS vehicle control, *L. murinus* KD6, or indole lactic acid (ILA). EAE was induced 4 days after the gavages began (n=18 vehicle (PBS); n=19 *L. murinus*; n=19 ILA; mean±SEM; two independent experiments). (E) Disease scores (\*\*\*\**p*-value<0.0001; two-way ANOVA; mean±SEM); (F) disease incidence rates (\**p*-value<0.05 or listed; Log-rank Mantel-Cox test; percentage); and (G) survival (\**p*-value<0.05, \*\**p*-value<0.01; Log-rank Mantel-Cox test; percentage) were tracked over 18 days post-immunization. (H) Mean disease severity of mice that developed disease (mean±SEM; each point represents a diseased mouse). (I) The proportion of mice with max disease score after 18 days. (J) Relative abundance of ILA-producing bacteria in control (n=576) and multiple sclerosis (MS) subjects (n=576) from iMSMS study^26^. \*\**p*-value<0.01; unpaired Wilcoxon test. (K) Relative abundance of ILA-producing bacteria in control (n=576), treated (n=367), and untreated (n=209) MS subjects. \*\**p*-value<0.01; \*\*\**p*-value<0.001; unpaired Wilcoxon tests. (L) Relative abundance of ILA-producing bacteria in MS subjects with mild, moderate, and severe disease. Mild: EDSS 0-3.5, no severe disability/highly ambulatory (n=414). Moderate: EDSS 4-6.5, severe disability while still ambulatory (n=140). Severe: EDSS 7-9.5, wheelchair bound (n=21). \*\**p*-value<0.01; \*\*\*\**p*-value<0.0001; unpaired Wilcoxon tests.

*Lactobacilli* are not the only source of ILA within the gut microbiota. ILA production has been observed in diverse bacterial species spanning multiple phyla^57^. We investigated the impact of βHB on the levels of ILA-producing microbes by re-analyzing published data for human *ex vivo* stool communities cultured with or without βHB for 24 hours^16^ (**Figure S7A-C**). We defined ILA-producing microbes as microbes with ≥4 fold-change over controls by mass spectrometry^57^. The relative abundance of ILA-producing microbes was significantly increased in response to βHB (**Figure S7A,B**). We hypothesized that βHB increased the abundance of ILA-producing microbes by selective growth promotion. Supporting this hypothesis, 12.5mM βHB treatment resulted in improved growth of *L. murinus* KD6 compared to a control bacteria, *B. adolescentis* BD1, as measured by shifted growth curves (**Figure S7C,D**) and decreased time to reach mid-exponential growth in monocultures grown in BHI media (**Figure S7E**). At higher concentrations (50mM) βHB inhibited growth of both *L. murinus* KD6 and *B. adolescentis* BD1 (**Figure S7C,D**); however, *L. murinus* growth was less inhibited than *B. adolescentis* as measured by a increased carrying capacity increased growth rate compared to the 0mM treatment (**Figure S7F,G**). Together, these data suggest that βHB shapes the microbiota composition to favor ILA-producing microbes by having a stronger growth inhibitory impact on non-ILA producing members of the microbiota communities.

We next tested the impact of *L. murinus* KD6 and ILA on EAE. We fed 6-8-week-old female CONV-R C57BL/6J mice a HFD for 3 days prior to delivering *L. murinus* KD6 (10^8^ CFUs) or ILA (20 mg/kg) by oral gavage every other day for the duration of the experiment. EAE was induced 4 days after the first gavage and disease was tracked for 18 days (**Figure 6E**). Disease scores over time were significantly lower in response to both *L. murinus* KD6 and ILA relative to vehicle controls (**Figure 6E**). Disease incidence was highest in the HFD group (**Figure 6F**). The probability of survival in the HFD group (28% by day 18) was significantly lower than the *L. murinus* (74%) and ILA (68%) treated groups (**Figure 6G**). 72% of the HFD controls reached a disease score >4.5, whereas the other two groups were mixed (**Figure 6H,I**). These data suggest that *L. murinus* protects mice during EAE due to the production of the anti-inflammatory metabolite ILA.

Notably, multiple studies support the clinical relevance of ILA where ILA is one of the most depleted serum metabolites in MS compared to controls^58,59^ and MS patients have a decreased abundance of ILA-producing gut bacteria^59^. We wanted to investigate these findings in an independent cohort of MS subjects (iMSMS) using our metric for ILA-producing microbes. ILA-producing microbes were significantly decreased in MS subjects compared to healthy controls (**Figure 6J**). This trend was driven by the untreated MS patients; treated subjects had similar levels of ILA-producing bacteria to controls (**Figure 6K**). ILA-producing bacteria were inversely associated with disease severity (**Figures 6L**). Taken together, these findings support the potential clinical relevance of our results in the EAE model.

## DISCUSSION

Herein, we provide evidence for a microbiome-dependent pathway through which a common dietary intervention protects from neurological disease. Building on prior studies in humans^6–8^ and mice^9–11^, we started by confirming that consumption of a KD rescues disease in the EAE mouse model. Prior to our work it remained unclear if the effects of a KD on EAE were microbiota-dependent. Our results suggest that assessing or even controlling for inter-individual variations in gut microbial community structure and function in future studies could potentially explain the variance in responsiveness to the KD observed in MS patients^6–8,60^ and between different mouse studies^9–11^.

Surprisingly, we discovered that oral delivery of a βHB-containing ketone ester is sufficient to mimic the protective effects of a KD. This finding, if replicated in humans, would suggest that βHB supplementation alone could be a viable therapeutic option instead of the full KD. The translational implications are profound, as KDs are limited by difficulties in maintaining such a strict diet long-term and could also have potential negative side effects^61^. Our identification of βHB as a key player provides a way to circumvent these barriers and provides a more general *proof-of-concept* for the ability to distill the activity of a complex diet down to a single bioactive molecule.

Our data from transgenic mice lacking the intestinal expression of *Hmgcs2* emphasizes the importance of studying the tissue-specific production of βHB and other diet-dependent metabolites. Surprisingly, the local production of βHB within the GI tract was required for the neuroprotective effect of a KD, despite no changes to circulating levels of βHB. These findings suggested that GI levels of βHB would be more informative of the effective dose of KE. Previously, we found that GI tissue levels of βHB were elevated to a similar extent by the KE supplementation as by a KD while serum and gut lumen levels were more elevated by a KD^16^. Together, these findings suggest that intestinal βHB levels may be an important measure of extent of protection mediated by a KE or KD. Future work will focus on the specific doses required to achieve a protective effect. These results add to the growing literature on interactions between the gut and neuroinflammation, showing how a host metabolite can indirectly signal to the brain via a downstream microbial effector.

We identified a specific *Lactobacillus* isolate, *L. murinus* KD6, and *Lactobacillus*-produced metabolite, ILA, that were each sufficient to rescue disease. Consistent with our results, supplementation with *L. murinus* reduced EAE disease severity in mice fed a high-salt diet^53^. Prior *in vitro* studies had shown that the *Lactobacillus-*produced metabolite ILA inhibits Th17 cells^53^ and our data shows that ILA is sufficient to protect from EAE. While the genes involved in bacterial production of ILA are well-characterized^52,54–56^, the generation of isogenic strains of *L. murinus* or other lactobacilli differing in the extent of ILA production would be valuable for dissecting the downstream host response in EAE or other disease models. These genetically engineered bacteria would enable more controlled clinical trials of MS patients given the potentially broad impacts of different probiotics for MS^62^. It is also possible that additional enzymes and pathways may impact ILA production in *L. murinus* KD6 that have not been characterized in other strains. There are multiple other potential mechanisms through which the gut microbiota can influence MS. Segmented filamentous bacteria (SFB or *Candidatus Arthromitus*) and an *Allobaculum* isolate exacerbate EAE due to Th17 cell activation^22,24^. In contrast, *Bacteroides fragilis* protects from disease due to elevated Foxp3+ regulatory T cells^63^. Inspection of our 16S-seq data did not reveal these specific EAE-associated species, suggesting that they do not contribute to our results. Future studies in gnotobiotic mice are warranted to test the interactions between these different bacterial taxa in promoting or suppressing disease in the context of different dietary inputs.

There are several limitations to our study deserving of follow-on work. βHB directly inhibits the growth of specific members of the microbiota^16^ but the mechanism of action and higher-level implications for microbial interactions remain unknown. While we opted to focus on the gut microbiome herein, the beneficial effects of diet for MS are likely also impacted by the direct effects of βHB on the host^33^, including inhibition of the inflammasome^34,35^. To complicate matters further, previous studies have highlighted the importance of the inflammasome in shaping the immunomodulatory potential of the microbiota, providing an indirect mechanism through which βHB could alter host-microbiome interactions^64–66^. Further, tryptophan metabolites including ILA signal through the aryl hydrocarbon receptor (AHR), which in turn influences Th17 responses^67–69^. Therefore, it will be important in future studies to investigate the role of host factors including the inflammasome and/or AHR in the mediation of the protective effect of a KD and KE supplementation.

Additionally, investigating other immunomodulatory effects of KD or KE shaped microbiota could provide additional insights and therapeutic avenues as Th17 cells are not the only immune responses contributing to MS and EAE. It will also be important to look at additional models of Th17 skewing such as with IL-23 and IL-1β, which have been implicated in EAE pathogenesis^70^. Diet-induced shifts in immune cell function could potentially contribute to the observed changes in the gut microbiota. It will be critical to conduct follow-on studies in humans and alternative mouse models of MS with additional dietary contexts and therapeutic interventions, to explore the translational relevance of ketone bodies, lactobacilli, and ILA. Finally, the mechanism of action through which ketone bodies impact gut bacterial growth and immunomodulatory potential remains underexplored and more work is needed to dissect the cellular and molecular pathways linking ILA and other tryptophan metabolites to disease.

Our work emphasizes the importance of considering both host and microbial metabolism, including key bioactive compounds like βHB and ILA. These results add to the growing literature on other diets that elevate βHB, including intermittent fasting and caloric restoration, which have shown some efficacy in MS patients and mouse models^9,71–74^. The pathway we have identified herein provides a common mechanism through which these diverse dietary interventions may impact MS. More broadly, our results set the stage to perform similar studies in other autoimmune diseases and for alternative dietary interventions, providing new insights into the complex mechanisms that link diet, the gut microbiome, and host immunity.

## Acknowledgments

We would like to acknowledge the UCSF Gnotobiotics Core Facility (gnotobiotics.ucsf.edu) and the Parnassus Flow Cytometry CoLab RRID:SCR_018206 supported in part by the DRC Grant NIH P30 DK063720. We also acknowledge the UCSF Quantitative Metabolite Analysis Center (QMAC) for their help quantifying ILA levels. We thank Dr. Michael Kattah for generously donating the VillinER/Cre mice. We thank BHB Therapeutics for providing the ketone ester. Funding was provided from the National Institutes of Health: R01DK114034, R01HL122593, R01AR074500, R01AT011117, P.J.T.; F32AI14745601, K99AI159227, M.A.; K08HL165106, V.U.; K08AR073930, D.O. and R.R.N; R01AG067333, J.C.N.; R01DK091538, R01AG069781, P.A.C. P.J.T is a Chan Zuckerberg Biohub-San Francisco Investigator and was a Nadia’s Gift Foundation Innovator supported, in part, by the Damon Runyon Cancer Research Foundation (DRR4216). Model figures were made using Biorender.com.

## Author contributions

Conceptualization: MA, PJT

Data curation: MA, VU, LR, RR, KT

Formal Analysis: MA, VU

Funding acquisition: RRN, MA, PJT, JCN

Investigation: MA, VU, QYA

Methodology: MA, VU, QYA, PP, PAC, RR, DO, LR, JAT, CW

Project administration: PJT

Resources: PJT, PAC, JCN

Software: VU

Supervision: PJT, RRN, JCN, PAC, AP

Validation: MA, VU

Visualization: MA, VU

Writing – original draft: MA

Writing – review & editing: PJT

## METHODS

### RESOURCE AVAILABILITY

#### Lead Contact

Further information and requests for resources and reagents should be directed to and will be fulfilled by the Lead Contact, Peter Turnbaugh.

#### Materials Availability

This study does not contain newly generated materials.

#### Data and Code Availability

All data is available in the main text, the supplemental tables, datasets, or deposited as listed below. This paper does not report original code. Any additional information required to reanalyze the data reported in this paper is available from the Lead Contact upon request. 16S-seq and *L. murinus* whole genome sequence deposited under Bioproject: PRJNA1032118

### EXPERIMENTAL MODEL DETAILS

#### Mice

All animal experiments were conducted under protocols (AN200526 and AN199820) approved by the UCSF Institutional Animal Care and Use Committee. Germ-free C57BL/6J mice were born at UCSF and CONV-R SPF C57BL6/J mice were purchased from Jackson Laboratory. Germ-free mice were maintained within the UCSF Gnotobiotic Core Facility. Mice age ranged from 8-16-week-old and mice were assigned to groups to achieve similar age distribution between groups. Mice were housed at temperatures ranging from 67 to 74 °F and humidity ranging from 30 to 70% light/dark cycle 12h/12h. No mice were involved in previous procedures before experiments were performed.

*Hmgcs2^WT^* (*Hmgcs2^fl/fl^ VillinER/Cre^-/-^*) or *Hmgcs2^ΔIEC^* (*Hmgcs2^fl/fl^ VillinER/Cre^+/-^*) mice on C57BL6/NJ substrain hybrid background were generated. *Hmgcs2^fl/fl^* mice were generated in collaboration with the inGenious Targeting Laboratory (Ronkonkoma, NY). Briefly, BF1 clone C57BL/6N mouse embryonic stem (ES) cells harboring an expression construct for FLP recombinase were electroporated with a donor vector targeting exon 2 of the *Hmgcs2* locus (NM_008256), along with Cas9 and synthetic guide RNAs. Targeting exon 2 results in a frame shift mutation and resulting nonsense (premature stop) encoding *Hmgcs2* transcript. After selection for neomycin resistance using G418, and confirmation of successful targeting using both Southern blots and PCR, targeted ES cells were were microinjected into Balb/c blastocysts, and chimeras with a high percentage black coat color were mated to C57BL/6N WT mice to generate Germline Neo Deleted mice (*Hmgcs2^fl/+^*). PCR and DNA sequencing were used to confirm (*i*) successful flanking of exon 2 with *loxP* sites, (ii) the loss of the neomycin resistance-encoding gene, and (*iii*) the absence of a FLP recombinase transgene in the progeny. . Transgenic mice harboring a tamoxifen-inducible Cre recombinase under the control of the villin-promoter (VillinER/Cre, C57BL/6J)^75^ were a gift from M. Kattah (UCSF). *Hmgcs2* was specifically and inducibly deleted from intestinal epithelial cells (IECs). These mice were made by crossing *Hmgcs2^fl/fl^* mice (shipped from the University of Minnesota) with *VillinER/Cre^+/-^* and then breeding mice to homozygosity. For experimental mice, *Hmgcs2^fl/fl^ VillinER/Cre^-/-^*mice were bred with *Hmgcs2^fl/fl^ VillinER/Cre^+/-^* mice to produce *Hmgcs2^WT^ - Hmgcs2^fl/fl^ VillinER/Cre^-/-^* and male mice with *Hmgcs2* specifically deleted from intestinal epithelial cells *Hmgcs2^ΔIEC^ - Hmgcs2^fl/fl^ VillinER/Cre^+/-^*. *Hmgcs2^WT^* and *Hmgcs2^ΔIEC^* mice were reared at UCSF.

*Hmgcs2^WT^* or *Hmgcs2^ΔIEC^* were induced with 5 daily i.p. injections of 100 μL of tamoxifen at a concentration of 10 mg/mL. Tamoxifen was prepared by dissolving 100 mg/mL of tamoxifen in 100% ethanol, incubating for 30 min at 56°C, vortexing, then diluting 1:10 in corn oil to a final concentration of 10 mg/mL. After tamoxifen injections, mice had a 5 day washout period before starting on their specific diets. Genotype of mice was confirmed with PCR where DNA was extracted from mouse tissue using Direct PCR (Thermo Fisher) according to the manufacturer’s instructions. Briefly, 150 μL of Direct PCR and 3 μL Proteinase K was added to the tissue then incubated in 56°C water bath overnight. 100 μL of the sample was then transferred to PCR strip tubes and incubated for 1 hour at 85°C. For PCR, 1 μL extracted DNA, 12.5 μL AmpliTaq Gold master mix (Thermo Fisher), 1 μL forward primer (1 µM final concentration), 1 μL reverse primer (1 µM final concentration) and 9.5 μL water was combined with the following primer sequences to detect Cre presence: Cre forward - 5’ TTC CCG CAG AAC CTG AAG ATG 3’; Cre reverse: - 5’ CCC CAG AAA TGC CAG ATT ACG 3’.

*Hmgcs2^fl/fl^* was homozygous and confirmed initially with the following primers: HMGCS2 forward - GTG AGT TCT GTG CCT GAC TG; HMGCS2 reverse - CAG AGC TGC AAG ATG AGT AAC TG. The following PCR cycle conditions were used for Cre detection (1. 95°C for 10 min; 2. 95°C for 30 sec; 3. 60°C for 30 sec; 4. 72°C for 30 sec; 5. Repeat steps 2-4 34x; 6. 72°C for 5 min; 7. 11°C hold) and for *Hmgcs2* detection (1. 95.0°C 10 min; 2. 95.0°C 5 sec; 3. 55.0°C 30 sec; 4. 72.0°C 10 sec; 5. Repeat steps 2-4, 29x; 6. 72.0°C 5 min; 7. 4.0°C hold). Only male mice were used for *Hmgcs2* experiments as in our initial experiments female mice on a KD did not develop clinical symptoms of disease in either *Hmgcs2^WT^* or *Hmgcs2^ΔIEC^*mice (*data not shown*). For non-transgenic mouse experiments both male and female mice were used for initial experiments and then female mice were used for later experiments to control for the potentially sexually dimorphic presentation of EAE^76^.

#### Diets

All diets were provided *ad libitum*. Semi-purified diets were purchased from Envigo, and the ingredients used in these diets are listed in **Table S1**. Before experimental diets started, mice were fed standard chow diet (Lab Diet 5058) in the SPF facility, while standard autoclaved chow diet (Lab Diet 5021) was used in the gnotobiotic facility. All diets used for gnotobiotic experiments were either autoclaved or irradiated to ensure sterility. For the design of the diets supplemented with KE: 10% C6X2-βHB KE was added at the expense of fat. C6X2-βHB is estimated to contain 8 kcal/g and is composed of fatty acids ester-linked to the ketone body β-hydroxybutyrate.

#### Antibiotic Water Treatments

Where specified mice were treated *ad libitum* with antibiotic water including 0.1 g/L ampicillin or a 4 antibiotic cocktail AVNM (1g/L <u>a</u>mpicillin, 0.5g/L <u>v</u>ancomycin, 1 g/L <u>n</u>eomycin, 1 g/L <u>m</u>etronidazole). Antibiotic water was prepared from autoclaved tap water with 0.2 μm filtering after the antibiotic(s) were added.

#### Experimental Autoimmune Encephalomyelitis (EAE) in C57BL/6J mice

EAE was induced with slight modification of previous descriptions^31^. An emulsion of a 100 mg/100 μL of pMOG_35–55_ (Tocris; cat# 25681) resuspended in sterile 1X phosphate buffered saline (PBS) and complete Freund’s adjuvant (CFA - 4 mg/mL: heat killed Mycobacterium tuberculosis (MTB) Fisher Scientific; cat# DF3114338 at 4 mg/mL in incomplete Freund’s adjuvant Millipore Sigma; cat# F5506) was prepared by passing through a three-way stopcock connected pair of 10 mL syringes for 5 minutes or until emulsion was stable by the water drop test. 100 μL of the MOG and CFA emulsion was injected into mice via subcutaneous injection. Directly following MOG injections, 500 ng of pertussis toxin (PTX) (Millipore Sigma; cat# P2708) in 100 μL of PBS was injected via i.p. injections. 500 ng PTX in 100 μL of PBS was again injected via i.p. the day following immunization. Mice were weighed and scored for disease daily for the duration of the experiment. EAE disease courses were tracked for 16-30 days for a combination of logistical and scientific rationales. To assess immune responses at peak disease some EAE experiments were tracked for shorter time periods.

EAE scoring was outlined as follows: 0=no obvious signs of disease; 0.5=partial tail weakness; 1=limp tail; 2=limp tail and hindlimb weakness; 2.5=limp tail and impairment in righting reflex; 3=one hindlimb paralyzed; 3.5=bilateral hindlimb paralysis; 4=bilateral hindlimb paralysis and forelimb paresis; 4.5=limp tail, complete hind leg and partial front leg paralysis, mouse is minimally moving around the cage but appears alert and feeding. Euthanasia required after the mouse scores 4.5 for 24 hours; 5=moribund: complete hind and partial front leg paralysis, no movement around the cage, mouse is not alert, has minimal movement in the front legs, or barely responds to contact; euthanasia as soon as possible.

#### Experimental Autoimmune Encephalomyelitis (EAE) in SJL mice

EAE was induced with slight modification of previous descriptions^38,39^. 9-week-old female SJL mice purchased from Jackson Labs were fed HFD, HFD-KE, or KD for 5 days prior to EAE induction. Mice that developed tail lesions (1 animal, in our case) were immediately euthanized and removed from the experiment given SJL mice with tail lesions do not develop EAE^39^. To induce EAE, an emulsion of a 100 ug/100 μL of PLP_139–151_ (Genscript; custom synthesis with >95% purity by HPLC and <10EU/mg endotoxin, sequence: HSLGKWLGHPDKF [unmodified]) resuspended in sterile 1X PBS and CFA (4 mg/mL: heat killed Mycobacterium tuberculosis (MTB) Fisher Scientific; cat# DF3114338 at 4 mg/mL in incomplete Freund’s adjuvant Millipore Sigma; cat# F5506) was prepared by passing through a three-way stopcock connected pair of 10 mL syringes until emulsion was stable by the water drop test. 100 μL of the PLP and CFA emulsion was injected into mice via subcutaneous injection. Directly following PLP injections, 200 ng of pertussis toxin (PTX) (Millipore Sigma; cat# P2708) in 50 μL of PBS was injected via i.p. injections. 200 ng PTX in 50 μL of PBS was again injected via i.p. the day following immunization.

Mice were weighed and scored for disease every day for the duration of the experiment. Mice that became singly housed due to death of cagemates were provided additional enrichment. SJL EAE was scored in accordance with the Hooke EAE rubric and modified Hooke rubric for recovery^77^.

#### Fecal Microbiota Transplants

Fecal donor mice were put on their respective diets for 1-3 weeks prior to fecal collection with 2 donor mice cohoused per group. Donor fecal samples were collected from mice the day preceding transplantation and then frozen at −80°C before the next day, being resuspended in 1:10 g/mL sterile pre-equilibrated PBS in an anaerobic Coy chamber. Each sample was vortexed for 1 min and left to settle for 2 min, and the supernatants were then used for gavage. 100 μL of each sample was administered by oral gavage into the mice. Recipient mice were put on respective diets 3 days before receiving the AVNM cocktail of antibiotics in drinking water for 7 days (prepared as described above). AVNM water was then removed and daily gavage of FMT material began 7 days preceding EAE induction, then continued for 7 days post EAE induction.

1. *L. murinus* Engraftment
2. *L. murinus* KD6 was incubated aerobically at 37°C overnight in 10 mL MRS media (Difco Lactobacilli MRS agar, BD Biosciences, cat# 288210). Cultures were centrifuged 10 min at 2500 rpm, washed with sterile 1X PBS, centrifuged 10 min at 2,500 rpm, and then resuspended in sterile 1X PBS for oral gavage. Mice were gavaged with 200 μl of *L. murinus* KD6 (10^8^ CFU) every other day for the duration of the experiment.

#### Indole Lactic Acid (ILA) Treatment

A solution of ILA (DL-Indole-3-lactic acid; Sigma; cat# I5508) in sterile 1X PBS was made by vortexing before and after a 10 min incubation at 56°C to fully dissolve ILA. Mice received a dose of 20 mg/kg ILA every other day via oral gavage as described previously^78,79^. Control HFD mice received oral gavages of 200 μl sterile 1X PBS (vehicle) every other day.

### METHOD DETAILS

#### Brain Leukocyte Isolation

Brain leukocytes were isolated with slight modifications of previously described techniques^80^. Brains were dissected out of the mice and put into 4 mL ice cold 1X PBS in 6 well plates on ice. Then, brains were cut into small pieces with scissors until small enough to pass through an 18 gauge needle several times in PBS to homogenize tissue. This tissue homogenate was then passed through a 70 μm filter and filters were then washed with 10 mL ice cold 1X PBS. Filtered homogenate was centrifuged at 2,000 rpm for 10 min and the supernatant was removed. Cells were resuspended in 5 mL of 30% Percoll (VWR; cat# 89428-524) and then overlaid on top of 5 mL 70% Percoll at room temperature. Percoll gradients were then centrifuged at 2,000 rpm at 25°C for 25 min, with no brakes (deceleration set to 0) and lowest acceleration (acceleration set to 1). Cells at the interface were collected, washed with 1X PBS, and prepared for flow cytometry analysis.

#### Spleen Cell Isolation

Spleen cells were prepped through gentle mashing with a syringe plunger then filtered through a 40 μm filter. Cells were pelleted by centrifugation (2,000 rpm 5 min) and then treated with 1 mL 1X red blood cell (RBC) Lysis Buffer (Biolegend; cat# NC9067514) to lyse red blood cells. Cells were pelleted again (2,000 rpm, 5 min) and then washed in 1X PBS.

#### Flow Cytometry

Immune cells were isolated from the brain and spleen as described above. Surface staining for lymphocytes was done in staining buffer (1X Hank’s Balanced Salt Solution (HBSS), Life Technologies; cat# 14170161) supplemented with 10 mM hydroxyethyl piperazineethanesulfonic acid (HEPES, Fisher Scientific; cat# NC0734307), 2 mM ethylenediaminetetraacetic acid (EDTA, Fisher Scientific; cat# AM9260G), and 0.5% (v/v) heat inactivated fetal bovine serum (Life Technologies; cat# 10438026) for 20 min at 4°C. Cells were then washed twice in supplemented 1X HBSS and analyzed via flow cytometry. The following antibodies were used: anti-CD3 (Clone: 17A2, Fluorophore: FITC, Fisher Scientific; cat# 501129303), anti-TCRβ (Clone: H57-597, Fluorophore: PE-CF594, Fisher Scientific; cat# BDB562841), anti-CD4 (Clone: GK1.5, Fluorophore: BV786, Fisher Scientific; cat# BDB563331), and live/dead staining was performed using aqua LIVE/DEAD Fixable Dead Cell Stain Kit (Fisher Scientific; cat# 501121526). Cells were stimulated with a cell stimulation cocktail (Fisher Scientific; cat# 501129036) containing PMA and ionomycin according to the manufacturer’s instructions and Golgi Plug (1 μl/sample) (Fisher Scientific; cat# BDB555029) for 4-6 hours at 37°C. Cells were surface stained, washed, and then fixed/permeabilized in 100 μl fixation and permeabilization buffer (Fisher Scientific; cat# BDB555028). Cells were washed twice in permeabilization/wash buffer (Fisher Scientific; cat# BDB555028) and then stained for intracellular cytokines with the following antibodies: anti-IFNγ (Clone: XMG1.2; Fluorophore: BV421; Fisher Scientific; cat# BDB563376) and anti-IL17a (Clone: ebio17B7; Fluorophore: PE-Cyanine7; Fisher Scientific; cat# 501129407). Cells were washed twice with permeabilization/wash buffer and then placed in a staining buffer for flow cytometry analysis. Cell population gating was done using isotype and single stain controls. Gating strategies and which figures they correspond to are outlined in **Figure S1**. The flow cytometry data were collected with a BD LSR Fortessa and analyzed with FlowJo software (version 10.6.1). All antibodies are included in the **Key Resources Table**.

#### Immunoblotting

Flash frozen liver and isolated colonic intestinal epithelial cells (IECs) were lysed in RIPA buffer. IECs were isolated from mouse colons using the following workflow. Colons were splayed longitudinally with mucus removed and cleaned. Colons were then stored in complete RPMI - 10% heat inactivated fetal bovine serum (Life technologies; cat# 10438026), 100 units per mL penicillin and streptomycin (Thermo Fisher; cat# 15070063), glutamax (Gibco; cat# 35050061), sodium pyruvate (Life Technologies; cat# 11360070), HEPES (Fisher Scientific; cat# NC0734307), and nonessential amino acids (Life Technologies; cat# 11140050) on ice. Media was removed by filtering through a 100 μM filter, and remaining tissue incubated in 1X HBSS - without Ca2+ and Mg2+ containing 5 mM EDTA (Fisher Scientific; cat# AM9260G) and 1 mM DL-Dithiothreitol (DTT; VWR; cat# 97063-760) for 45 min at 37°C on a shaker at 200 rpm. Samples were filtered through a 100 μM filter and IECs were collected from the flow-through. Immunoblotting with an Hmgcs2 specific antibody (rabbit anti-mHMGCS2 (Santa Cruz Biotechnology Inc.; cat# sc-393256) at a 1:2000 dilution and GAPDH (Prointech; mouse anti-mGAPDH; cat# 60004) at a 1:10,000 dilution was performed. Secondary antibodies conjugated to horseradish peroxidase were goat anti-rabbit IgG (Southern Biotech; cat# 4030-05); 1:50000 for Hmgcs2 and Goat Anti-Mouse IgG (Abcam; cat# Ab6708); 1:2000 for anti-mGAPDH.

#### β-hydroxybutyrate (βHB) Quantitation

For measurements of circulating βHB, blood was obtained from mice immediately following euthanasia. Serum was collected with a serum separator (BD Microtainer cat# 365978), centrifuged at 13,000 rpm for 10 min at 4°C, and frozen at −80°C until use. IECs were isolated as described in the <u>Immunoblotting</u> section. Serum and IECs were extracted as previously described^81,82^. Briefly, samples were extracted with 1 mL (IECs) or 80 μL (serum) of ice cold methanol [80% v/v], followed by homogenization. After centrifugation, the supernatants were dried in vacuum and reconstituted in 100 μL (IECs) or 50 μL (serum) of methanol (3% v/v). Quantitative analysis was performed with a Dionex UltiMate 3000 quaternary HPLC system connected to Exactive^TM^ Plus Orbitrap mass spectrometer (Thermo Fisher Scientific, Waltham, MA) with a Waters XSelect HSS T3 column (2.1 mm x 100 mm, 2.5 µm). Data were normalized by a ^13^C stable isotope labeled internal standard (Cambridge Isotope Laboratories, Inc., Tewksbury, MA). The results were quantified using a standard curve with concentrations ranging from 0.78 μM to 50 μM. For quantification of βHB during EAE experiments, βHB was quantified using the Precision Xtra blood glucose and ketone monitoring system (Abbott; cat# 98814-65) using blood β-Ketone test strips (Abbott; cat# ART07249) in *ad libitum* fed mice in the morning (in the 1st 6 hours of the 12 hour light cycle). Briefly, a drop of tail blood was applied to the β-Ketone test strip inserted in the Precision Xtra blood.

### Stable *In Vitro* Communities (SICs)

SICs were derived from HFD, HFD supplemented with a βHB ketone ester (HFD-KE), or a ketogenic diet (KD) mouse stool sample collected 7 days post EAE induction (1 donor/group, 4-5 independent SICs/donor sample). Incubations were performed anaerobically, as described^45^. Briefly, 100 mg fecal matter was resuspended in 1 mL brain heart infusion media supplemented with L-cysteine (0.05% w/v), hemin (5 μg/mL), and vitamin K (1 μg/mL) (BHI^CHV^) with 15% glycerol. Samples were vortexed to suspend the fecal pellets. Sample debris was allowed to settle for 5 min and 500 μL of the supernatant was transferred to a fresh tube. In a 96 well plate, 195 μL of BHI^CHV^ media and 5 μL of each fecal slurry was added to each well. Communities were grown for 48 hours at 37°C in an anaerobic chamber (Coy Laboratory Products, 2%–5% H_2_, 20% CO_2_, balance N_2_). Communities were passaged a total of 3 times (initial incubation with 2 additional passages) by adding 5 μL of the previous culture to 195 μL of fresh BHI^CHV^ media and growing an additional 48 hours. Cell pellets from the third passage were then used to extract DNA for 16S rRNA gene sequencing. Cell free supernatants (CFS) were collected by centrifuging samples for 10 min at 2,500 rpm and then passing the resulting supernatant through a 0.2 μm filter for use in our Th17 skewing assay.

#### Bacterial Strain Isolation, Culturing, and Identification

To isolate individual strains from our SICs we plated on MRS (Difco Lactobacilli MRS agar, BD Biosciences, cat# 288210) plates at 37°C in an anaerobic chamber (Coy Laboratory Products, 2%–5% H_2_, 20% CO_2_, balance N_2_) for 48 hours and selected for individual colonies with streak purification of colonies with distinct morphologies. *L. murinus* KD6 was isolated from a KD-SIC (KD-SIC replicate 4 - from **Figure 4A**) on MRS media plates. The *L. murinus* KD6 isolate was identified as *Lactobacillus murinus/animalis* by 16S rRNA gene sequencing amplifying between V1 and V9 with bi-directional sequencing (Azenta Life Sciences) and annotation using NCBI BLASTN with the rRNA_typestrains/16S_ribosomal_RNA database. Species-level annotation to *Lactobacillus murinus* was independently validated using a MALDI Biotyper Sirius (Bruker) according to manufacturer’s instructions with direct transfer, extended direct transfer, and extraction sample preparation methods according to manufacturer’s instructions. Briefly, for direct transfer for each sample a smear of an isolated colony was applied as a thin film to the MALDI target plate (MBT Biotarget 96 US IVD; cat# 1840380). Then, 1 µL of matrix solution (10 mg/mL of α-cyano-4-hydroxy-cinnamic acid (HCCA) in Bruker’s solvent (50% acetonitrile, 47.5% water, 2.5% trifluoroacetic acid)) was added on top of smear and dried at room temperature. For extended direct transfer, the same procedure was performed as the direct but with an additional step of adding 1 µL of 70% formic acid and air drying at room temperature before matrix application. For extraction sample preparation, using an inoculation loop, a single colony was transferred to 300 µL of HPLC-grade water and mixed until the material was in suspension. Then 900 µL of 100% ethanol was added and mixed by vortexing. Samples were pelleted with a 2 min 15,000 rpm centrifugation and the supernatant was removed. Samples were pelleted again with a 2 min 15,000 rpm centrifugation and residual ethanol was removed. Samples were then air dried at room temperature for at least 5 min. Then, 25 µL of 70% aqueous formic acid was added and vortexed to resuspend the pellet. 25 µL of 100% acetonitrile was then added and mixed by vortexing. Samples were then centrifuged 2 min 15,000 rpm and 1 µL of the supernatant was applied to the MALDI target plate. A BTS (*Escherichia coli*) was used as a calibration standard.

#### DNA Extraction and Whole Genome Sequencing of *L. murinus* KD6

DNA extraction and whole genome sequencing with Illumina and Nanopore sequencing was performed by Seqcenter (https://www.seqcenter.com/).

##### DNA extraction

ZymoBIOMICS DNA Miniprep Kit. A cell pellet (10^9^ CFU) of *L. murinus* KD6 was resuspended in 750 µL of lysis solution. Cells suspended in lysis solution were transferred into the ZR BashingBeadTM Lysis Tubes and mechanically lysed using the MP FastPrep-24 lysis system with 1 minute of lysis at maximum speed and 5 minutes of rest for 5 cycles. Samples were then centrifuged at 10,000 rcf for 1 minute. 400 µL of supernatant was transferred from the ZR BashingBead Lysis Tube to a Zymo-Spin III-F Filter and centrifuged at 8,000 rcf for 1 minute. 1,200 µL of ZymoBIOMICS DNA Binding Buffer was added to the effluent and mixed via pipetting. 800 µL of this solution was transferred to a Zymo-Spin IICR Column and centrifuged at 10,000 rcf for 1 minute. This step was repeated until all material was loaded onto the Zymo-SpinTM IICR Column. DNA bound to the Zymo-Spin IICR Column was washed 3 times with 400 µL and 700 µL of ZymoBIOMICS DNA Wash Buffer 1 and then 200 µL of ZymoBIOMICS DNA Wash Buffer 2 with a 1-minute spin down at 10,000 rcf for each, respectively. Washed DNA was eluted using 75 µL of ZymoBIOMICS DNase/RNase Free Water following a 5-minute incubation at room temperature and a 1-minute spin down at 10,000 rcf. The Zymo-Spin III-HRC Filter was prepared using 600 µL of the ZymoBIOMICS HRC Prep Solution and a centrifugation at 8,000 rcf for 3 minutes. Eluted DNA was then purified by running the effluent through the prepared Zymo-SpinTM III-HRC Filter. Final DNA concentrations were determined via Qubit.

##### Nanopore

Sample libraries were prepared using the PCR-free Oxford Nanopore Technologies (ONT) Ligation Sequencing Kit (SQK-NBD114.24) with the NEBNext Companion Module (E7180L) to manufacturer’s specifications. No additional DNA fragmentation or size selection was performed. Nanopore sequencing was performed on an Oxford Nanopore a MinION Mk1B sequencer or a GridION sequencer using R10.4.1 flow cells in one or more multiplexed shared-flow-cell runs. Run design utilized the 400 bps sequencing mode with a minimum read length of 200bp. Adaptive sampling was not enabled. Guppy1 (v6.4.6) was used for super-accurate basecalling (SUP), demultiplexing, and adapter removal. No quality trimming has been performed.

##### Illumina

Illumina sequencing libraries were prepared using the tagmentation-based and PCR-based Illumina DNA Prep kit and custom IDT 10bp unique dual indices (UDI) with a target insert size of 320 bp. No additional DNA fragmentation or size selection steps were performed. Illumina sequencing was performed on an Illumina NovaSeq 6000 sequencer in one or more multiplexed shared-flow-cell runs, producing 2×151bp paired-end reads. Demultiplexing, quality control and adapter trimming was performed with bcl-convert1 (v4.1.5).

##### Illumina and Nanopore Combo Assembly and Annotation

Porechop^83^ (version 0.2.4; default parameters) was used to trim residual adapter sequence from the Oxford Nanopore Technology (ONT) reads that may have been missed during basecalling and demultiplexing. De novo genome assemblies were generated from the Oxford Nanopore Technologies (ONT) read data with Flye^84^ (version 2.9.2) under the nano-hq (ONT high-quality reads) model. Additional Flye options initiate the assembly by first using reads longer than an estimated N50 based on a genome size of 6Mbp. Subsequent polishing used the Illumina read data with Pilon^85^ (version 2.5.1) under default parameters. To reduce erroneous assembly artifacts caused by low quality nanopore reads, long read contigs with an average short read coverage of 15x or less were removed from the assembly.

Assembled contigs were evaluated for circularization via circulator^86^ (version 1.5.5) using the ONT long reads. Assembly annotation was then performed with Bakta^87^ (version 1.8.1) using the Bakta v5 database. Finally, assembly statistics were recorded with QUAST^88^ (version 5.2.0).

#### ILA Detection via Liquid Chromatography Mass Spectrometry (LCMS)

##### Bacterial cell free supernatants

Cell free supernatant was obtained from *L. murinus* KD6 and *L. reuteri* strain ATCC23272 24 hour anaerobic cultures in BHI^CHV^. Cell free supernatants (CFS) were collected where cultures were pelleted via centrifugation at 3,500 rpm, for 10 minutes and then the resulting supernatant was passed through a 0.22 µm filter and stored at −80°C.

##### CFS extraction for LCMS

To assess ILA production CFS were extracted with acetonitrile (ACN) and methanol (MeOH) where 250 µL of CFS was aliquoted and 1 mL of a 50:50 ACN:MeOH was added. 100 pg/µL of a stable isotope labeled amino acid mixture for mass spec (Sigma Aldrich; cat# 909653) and 2-Amino-3-bromo-5-methylbenzoic acid was added to each sample and vortexed briefly. Samples were then incubated on ice for 15 minutes then centrifuged at max speed at 4°C on a tabletop centrifuge and resulting supernatants were transferred to a new tube. Samples were then dried with a speedvac and resuspended in 200 µL LCMS grade water.

##### Targeted Metabolomics

Samples were analyzed using a SCIEX ExionLC UPLC in series with SCIEX 7500 QTrap. 5 ml of each sample were injected and separated using Phenomenex’s Kinetex 2.6 mm F5 100 Å LC column 2.1 x 150 mm (cat # 00F-4723-AN). The mobile phase scheme consisted of A: H_2_O + 0.01% formic acid and B: Acetonitrile + 0.01% formic acid. At a constant flow rate of 200 ml/min, metabolites were separated using the following gradient: 0.2 % B (initial), 0.2% B (2 min), 5% B (5 min), 25% B (11 min), 98% B (13 min), 98% B (16 min), 0.2% B (16.1 min), and 0.2% B (20 min). Data was acquired in positive mode with optimized MRM pair determined from a commercial standard. The following source parameters were used: Ion source gas 1 = 45, Ion source gas 2 = 70, CUR = 40, CAD = 9, TEM = 375, Ion Source = 2000, and pause time = 5 ms. The following MRM pairs were used: 3-Indolelactic acid - Q1 = 206.1, Q3 = 188.1, CE = 20, ^15^N_2_-Tryptophan - Q1 = 207.2, Q3 = 147.1, CE = 20. EP and CXP values were held at 10 and 15 and dwell time = 5 ms for both MRM pairs. Acquired data was analyzed using SCIEX OS.

#### Th17 Skewing Assay

Th17 skewing was performed as described^50^. RBC lysed mouse splenocytes from a male C57BL/6J CONV-R 17-week-old mouse were filtered through a 40 μm filter and used for T cell isolation. Naive CD4+ T cells were isolated via Dynabeads untouched mouse CD4 isolation kit (ThermoFisher) according to kit specifications. In a 96 well plate pre-coated with anti-CD3 (5 μg/mL, overnight 37°C), equal cell numbers were plated and were treated with CFS or media controls at 5% v/v. At the same time, Th17 skewing conditions were supplied: 10 μg/mL anti-CD28, 0.3 ng/mL TGFbeta, 20 ng/mL IL-6, 2 ng/mL anti-IFNgamma, and 2 ng/mL anti-IL-4^51^. Isolated CD4+ T cells were developed in Th17 skewing conditions with bacterial CFS present for 4 days at 37°C and then restimulated with a cell stimulation cocktail (Fisher Scientific) containing PMA and ionomycin, according to the manufacturer’s instructions. Cells were restimulated at 37°C overnight and then supernatants were harvested for IL-17a quantification via ELISA. We utilized the mouse IL-17a (homodimer) ELISA (ThermoFisher) according to the kit instructions, where 100 μL of cell culture media was loaded onto the ELISA. Absorbance for ELISA was measured at 450 nm and the blank background signal was subtracted.

#### DNA Extraction

Mouse fecal samples (30-60 mg of preweighed material) were homogenized with bead beating for 5 min (Mini-Beadbeater-96, BioSpec) using the ZR BashingBead lysis matrix containing 0.1 and 0.5 mm beads (ZR-96 BashingBead Lysis Rack, Zymo Research; cat# S6012-50) and the lysis solution provided in the ZymoBIOMICS 96 MagBead DNA Kit (Zymo Research; cat# D4302). Bead beating was repeated for a total of three 5 min rounds with 5 min incubation at room temperature in between. Samples were then incubated at 65°C for 10 min before centrifuging for 5 min at 4,000 rpm. The supernatant was transferred to 1 mL deep-well plates and the DNA was then purified using the ZymoBIOMICS 96 MagBead DNA Kit according to the manufacturer’s instructions.

#### 16S rRNA Gene Sequencing

The V4 region of the 16S rRNA gene was amplified using primers 515F and 806R (see **Key Resources Table**). The reaction mix was: 0.45 µL DMSO for PCR (Sigma; cat# D8418-50mL), 0.0045 µL SYBR Green I (Sigma; cat# S9430) 10x diluted in DMSO to 1000x, KAPA HiFi PCR kit (1.8 µL 5x KAPA HiFi Buffer, 0.27 µL 10 mM dNTPs, 0.18 µL KAPA HiFi polymerase; cat# KK2502), 0.045 µL of each of the amplification primers 515F and 806R (final concentration 1 µM), 6.2055 µL Nuclease-free H2O (Life Tech; cat# 0977-023) and 1 µL DNA. We used a BioRad CFX 384 real-time PCR instrument with four serial 10-fold dilutions of extracted DNA template (10-100 ng/µL). Amplification was as follows: 5 min at 95°C, 20x (20 sec at 98°C, 15 sec at 55°C, 60 sec at 72°C), and hold at 4°C. Individual sample dilutions with amplification in the exponential phase (non-plateaued) were selected for subsequent indexing PCR using dual GoLay index primers^89^. For indexing KAPA HiFi PCR kit was used with, 4 µL 5x KAPA HiFi Buffer, 0.6 µL 10 mM dNTPs, 1 µL DMSO, 0.4 µL KAPA HiFi polymerase, 4 µL of indexing primer^89^, and 10 µL of 100 fold dilution of a non-plateaued primary PCR reaction was used with the same amplification as above. Amplicons were quantified with PicoGreen (Quant-It dsDNA, Life Technologies; cat# P11496) according to manufacturer’s instructions and pooled at equimolar concentrations. Aliquots of the pools were then column (MinElute PCR Purification Kit, QIAGEN; cat# 28004) and gel purified (QIAquick Gel Extraction Kit, QIAGEN; cat# 28704). Libraries were then quantified using KAPA Library Quantification Kit for Illumina Platforms (KAPA; cat# KK4824) according to manufacturer’s instructions and sequenced with an Illumina MiSeqV3 using 15% PhiX spike in. All sequencing was paired end, with 270 bp fragments, index length of 8 bp, mean library size 435 bp.

#### 16S rRNA Gene Analysis

Primer sequences and adapters were trimmed using the cutadapt plugin in QIIME2^90^, and were truncated to 220 base pairs for forward sequences and 150 bp for reverse sequences. Sequences then underwent quality filtering, denoising, and chimera filtering utilizing DADA2 (dada2 v 1.18.0)^91^, using default parameters (QIIME command PairedEndFastqManifestPhred33). Taxonomy was assigned to amplicon sequence variants (ASVs) using the SILVA v138 database^92^. Sequence variants were filtered such that they were present in >3 samples with ≥10 reads using MicrobeR (version 0.3.2)^93^. Data was then transformed using a Centered log2-ratio (CLR) with a CZM pseudocount replacement by zCompositions version 1.4.0-1^94^. CLR data was transformed into a Euclidean distance matrix using the dist command from the stats package in R (version 4.2.1). PERMANOVA testing was conducted on this distance matrix using adonis2 and Shannon diversity was calculated using the diversity command from vegan (version 2.6.4)^95^. Filtered counts were used in differential abundance. Differential abundance testing of ASVs was completed using ALDEx2^96^. A phylogenetic tree of statistically significant features was created using the ggtree package version 3.6.2^97^.

#### *Ex vivo* Incubation and Sequencing of Human Stool Samples Analysis

We re-analyzed data from Ang et al. 2020^16^ to assess changes in ILA-producing microbes in human *ex vivo* communities exposed to beta-hydroxybutyrate. *Ex vivo* communities were generated from human fecal samples and incubated for 24 hours in BHI^CHV^ media in the presence or absence of 12.5 mM beta-hydroxybutyrate in quadruplet, followed by DNA extraction and 16S rRNA sequencing as described in^16^. Demultiplexed sequences were processed with a 16S rRNA gene analysis pipeline and mapped to species abundances with qiime2R and phyloseq as described in^98^. Relative abundances of strains with documented ILA production (≥4 fold-change over controls)^57^, were binned and compared using the mixed-effects model Abundance ∼ Treatment + 1|Donor using the nlme package^99^.

#### ILA-producer Analysis in iMSMS

We downloaded patient metadata and the ASV read table from the iMSMS consortium^26^. We reassigned taxonomy based on SILVA SSU database 138 in RStudio and normalized as proportions. Using these data, relative abundances of strains with documented ILA production (≥4 fold-change over controls)^57^ were assessed. ILA-producing species proportions were summed and converted to percentage of total ASVs. We binned multiple sclerosis severity based on EDSS, a standardized disease scoring rubric for multiple sclerosis, with EDSS 0-3.5 classified as mild, EDSS 4-6.5 (severe disability, highly ambulatory) classified as moderate, and EDSS 7-9.5 (wheelchair bound) classified as severe.

#### Growth Curves of *L. murinus* KD6 and *B. adolescentis* BD1

A 96 well plate was filled with 150 μL of BHI supplemented with L-cysteine (0.05% w/v) and resazurin (0.0001%, w/v) (BHI^CR^) and 7 colonies of *L. murinus* KD6 and *B. adolescentis* BD1 were grown overnight. 1.5 μL of these turbid cultures was used as starter cultures into 150 μL BHI^CR^ with three concentrations of βHB (Sigma 166898) - 0 mM, 12.5 mM, and 50 mM from a 0.22 uM sterile filtered 1M solution of βHB made in BHI^CR^. This was then loaded onto the EON plate reader, set with continuous shaking and OD every 15 minutes. For background blank subtraction, multiple replicates of 0mM, 12.5mM, and 50mM βHB concentrations in BHI^CR^ were used. For 1 biological replicate of the *B. adolescentis* BD1 50mM condition, growth metrics could not be measured due to a poor model fit as defined as [(carrying capacity < 0.1 & growth rate > 0.5) or carrying capacity > 3 or time to mid exponential < 0, Sample] using the in growth curver tool^100^ (R package growthcurver 0.3.1).

#### Quantification and Statistical Analysis

Statistical tests, the software used, the number of replicates, definitions of center, and dispersion/precision measures are specified in the figure legends or on the plots themselves where possible. Tests were two-tailed unless otherwise noted. Statistical analyses were performed using GraphPad Prism software (Version 8) QIIME2 and R. Categorical data across 3 or more experimental groups was analyzed using ANOVA Kruskal-Wallis with Dunn’s multiple comparison test and one-way ANOVA with Dunnett’s multiple comparisons test for comparisons to BHI^CHV^ control in Th17 skewing assay. Unpaired Welch’s *t*-tests were used for comparisons of two groups. Longitudinal data was analyzed using two-way ANOVA. For 16S rRNA gene analysis, PERMANOVA testing was conducted on this distance matrix using adonis2. For correlations, Spearman correlations were performed with the exception of the correlation between ΔCLR HFD-KE vs. HFD and ΔCLR KD vs. HFD which is a Pearson correlation due to the type of data being matched between axes.

## SUPPLEMENTAL FIGURES AND FIGURE LEGENDS

**Figure S1.**
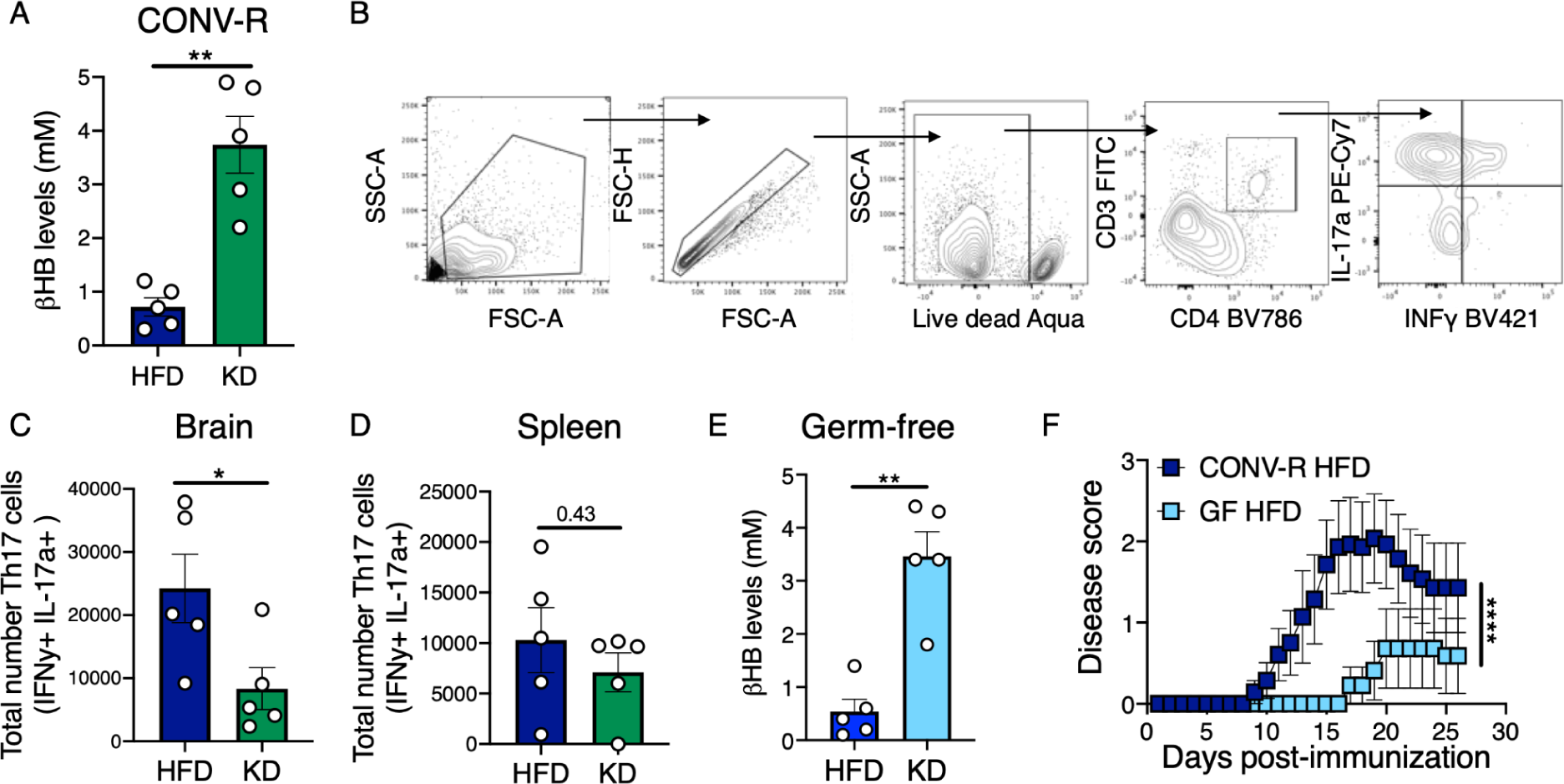
βHB levels are increased on a KD in both CONV-R and GF EAE mice and flow cytometry gating strategies. Related to Figures 1 and 3. (A) Circulating βHB levels in HFD or KD fed CONV-R EAE mice 16 days post-immunization from experimental mice in Figure 1A-H (\*\**p*-value<0.01; Welch’s *t*-tests; mean±SEM). (B) Flow cytometry gating strategies. Lymphocytes were gating on FCS-A and SSC-A, then gated on singlets with FCS-A and FCS-H. Cells were then gated on either (top) live cells, CD3+ CD4+ cells, then on IL-17a and IFNγ. (C-D) Total Th17 cell numbers in the (C) brain and (D) spleen of HFD or KD EAE mice (related to Figure 1E-H). Th17 cells were gated on single, live, CD3+ CD4+, and the number of IL-17a+ and IFNγ+ are quantified (\**p*-value<0.05, listed; Welch’s *t*-tests; mean±SEM). (E) Circulating βHB levels in HFD or KD fed germ-free EAE mice 32 days post-immunization from experimental mice in Figure 1I-J from one independent experiment (\*\**p*-value<0.01; Welch’s *t*-tests; mean±SEM). (F) Replotting of EAE disease scores over 16 days post immunization of HFD fed CONV-R versus GF mice from Figure 1A and I (\*\*\*\**p*-value<0.0001; two-way ANOVA; mean±SEM).

**Figure S2.**
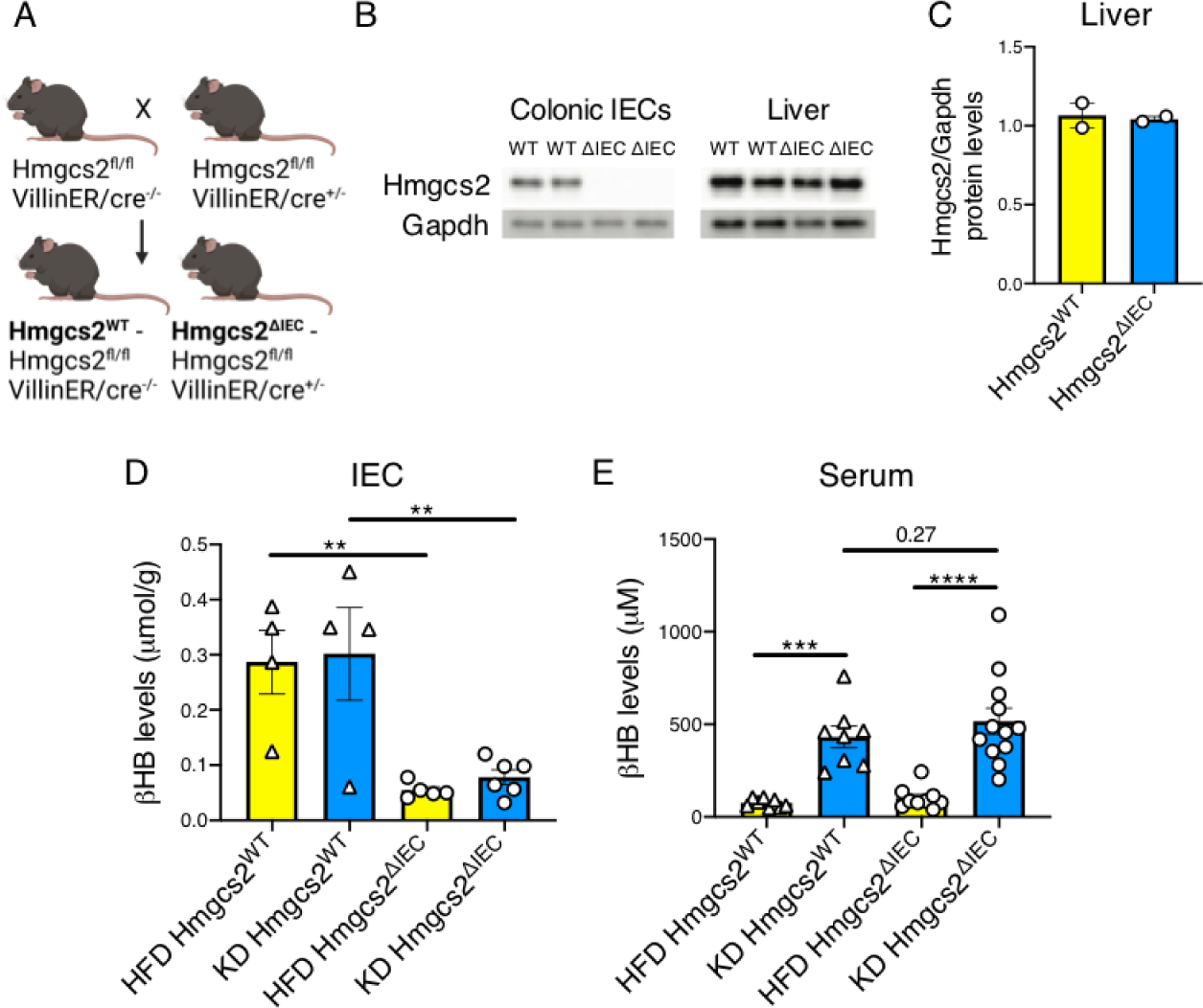
*Hmgcs2^ΔIEC^* design and validation. Related to Figure 3. (A) *Hmgcs2* was specifically and inducibly deleted from intestinal epithelial cells (IECs). These mice were made by crossing *Hmgcs2^fl/fl^* mice with *VillinER/Cre^+/-^* and then breeding mice to homozygosity. For experimental mice, *Hmgcs2^fl/fl^ VillinER/Cre^-/-^* mice were bred with *Hmgcs2^fl/fl^ VillinER/Cre^+/-^* mice to produce WT controls (*Hmgcs2^WT^ - Hmgcs2^fl/fl^ VillinER/Cre^-/-^*) and mice with *Hmgcs2* specifically deleted from intestinal epithelial cells (*Hmgcs2^ΔIEC^ - Hmgcs2^fl/fl^ VillinER/Cre^+/-^*). (B) To validate the model, we induced the deletion of *Hmgcs2* with daily 100 µl intraperitoneal injections of 10 mg/ml tamoxifen for 5 days and then 10 days later assessed the deletion of *Hmgcs2* in IECs and liver as well as the impact on βHB levels both in the IEC compartment and serum of these mice. Western blot of *Hmgcs2* and Gapdh from *Hmgcs2^W^*^T^ and *Hmgcs2^ΔIEC^* mice in colonic IECs and liver lysates. (C) Quantification of liver *Hmgcs2* levels from (B) relative to Gapdh loading control as quantified with ImageJ. (D-E) βHB levels were measured via mass spectrometry in (D) colonic IECs or (E) serum from male and female *Hmgcs2^WT^* (triangles) and *Hmgcs2^ΔIEC^*(circles) treated with 5 daily treatments with 100 µl intraperitoneal injections of 10 mg/ml tamoxifen and a week wash out period then either a HFD (yellow) or KD (blue) for 3 weeks (\*\**p-*value<0.01, \*\*\**p-*value<0.001, \*\*\*\**p-*value<0.0001, listed; one-way ANOVA with Holm-Sidak’s multiple comparisons test; mean±SEM; n=4-6 IECs; n=8-12 serum).

**Figure S3.**
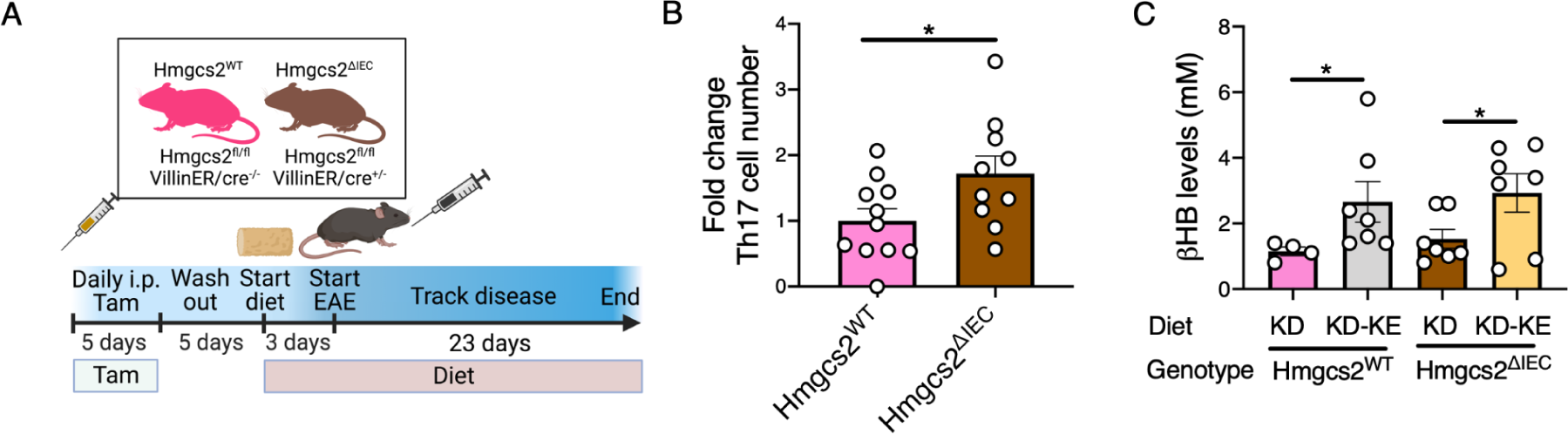
βHB levels are increased with KD-KE supplementation but not altered between *Hmgcs2^ΔIEC^* and *Hmgcs2^WT^* mice during EAE. Related to Figure 3. (A) Experimental design for experimental autoimmune encephalomyelitis (EAE) experiments with *Hmgcs2^WT^*and *Hmgcs2^ΔIEC^* mice. Mice were treated with 5 daily treatments with 100 µl intraperitoneal injections of 10 mg/ml tamoxifen and a 5 day wash out period before being put on a KD (for Figure 3A-H) or a KD or KD-KE (for Figure 3I-L) for 3 days. Then EAE was induced and disease was tracked. (B) Fold-change in the number of Th17 cells in the brain of *Hmgcs2^WT^* and *Hmgcs2^ΔIEC^*EAE mice (related to Figure 3F-G). Th17 cells were gated on single, live, CD3+ CD4+, and the number of IL-17a+ and IFNγ+ are quantified (\**p*-value<0.05, Welch’s *t*-test; mean±SEM). (C) βHB levels 7 days post-immunization in the serum of *Hmgcs2^WT^* and *Hmgcs2^ΔIEC^* fed a KD or KD-KE (\**p*-value<0.05; one-tailed Welch’s *t*-tests; mean±SEM).

**Figure S4.**
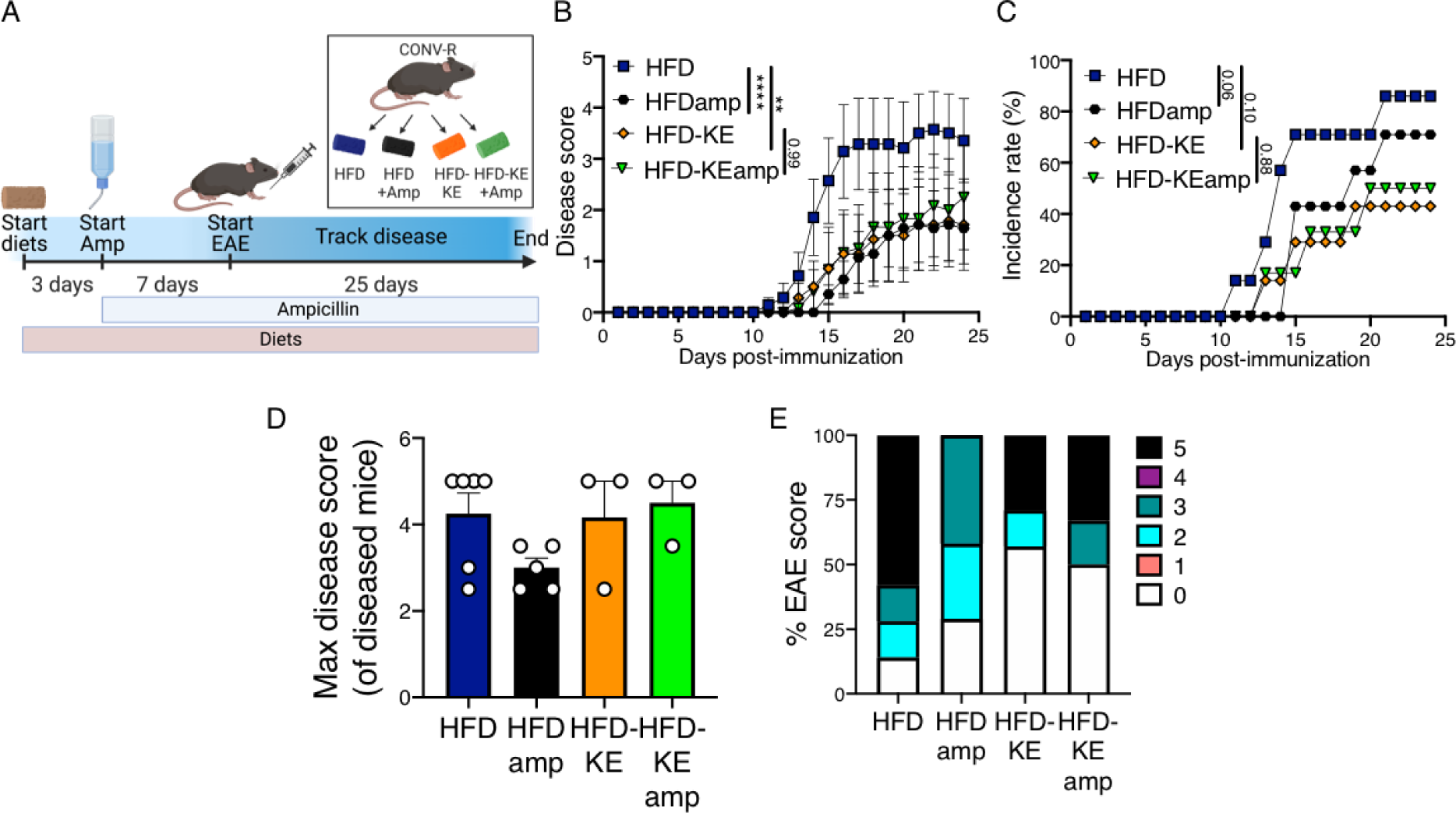
The impact of antibiotic treatment on disease is diet-dependent. Related to Figure 4. (A-F) Conventionally raised (CONV-R) specific pathogen free female Jackson mice were fed a high-fat diet (HFD) or a HFD supplemented with a βHB ketone ester (HFD-KE) for 3 days prior treatment with or without ampicillin water 7 days prior to EAE disease induction (n=7 HFD, HFD+amp, HFD-KE; n=6 HFD-KE+amp). (B) Disease scores (**p<0.01; ****p<0.0001; listed; two-way ANOVA; mean±SEM) and (C) disease incidence (*p*-value listed; Log-rank Mantel-Cox test; percentage) were tracked over days post-immunization. (D) Mean disease severity of mice that developed disease (mean±SEM; each point represents a disease mouse). (E) The percentage of mice with max disease score after 25 days.

**Figure S5.**
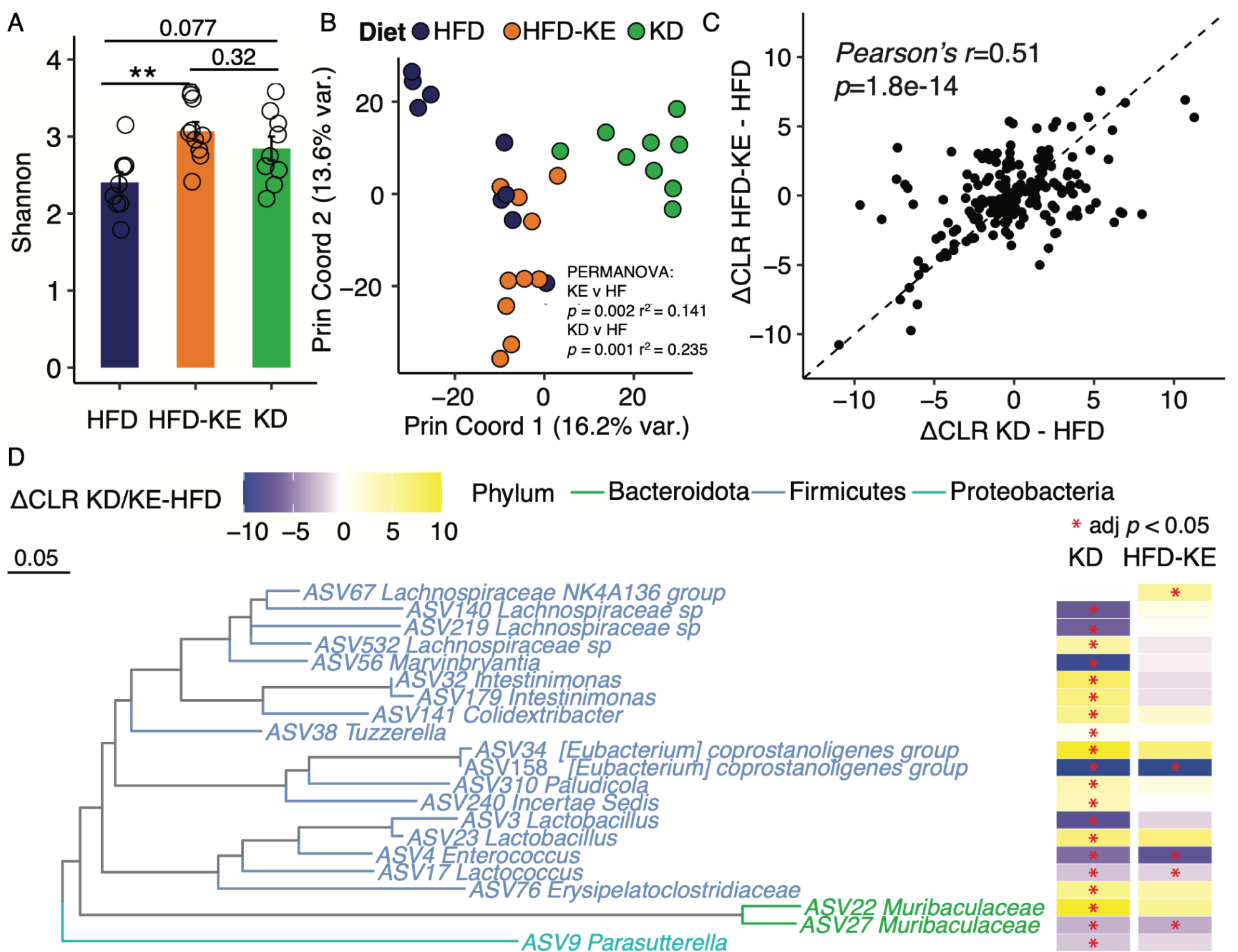
Ketogenesis alters the gut microbiotas of diseased mice. Related to Figure 2. (A-D) 16S rRNA gene sequencing was performed on a high-fat diet (HFD), HFD supplemented with ketone ester (HFD-KE), and ketogenic diet (KD) fed C57BL/6J EAE mice day 7 post-immunization (n=9 HFD; n=10 HFD-KE and KD; same mice as in Figure 2C-D). (A) Shannon diversity in HFD-KE, HFD, and KD fed EAE mice (*p*-values listed; Kruskal-Wallis with Dunn’s test). (B) Principal coordinate analysis (PCA) of Euclidean distances of centered log ratio (CLR) transformed amplicon sequence variants (ASVs; *p*-values and r^2^ values, PERMANOVA). (C) Correlation between the change in the CLR of ASVs of HFD-KE compared to HFD (ΔCLR HFD-KE vs. HFD) and KD compared to HFD (ΔCLR KD vs. HFD) which revealed a significant correlation (Pearson’s correlation rho and *p*-value shown) (D) Phylogenetic tree and change in center log ratio (ΔCLR) for significantly altered amplicon sequence variants (ASVs) between HFD versus KD and HFD versus HFD-KE. (right of panel; * denotes adjusted *p*-value<0.05)

**Figure S6.**
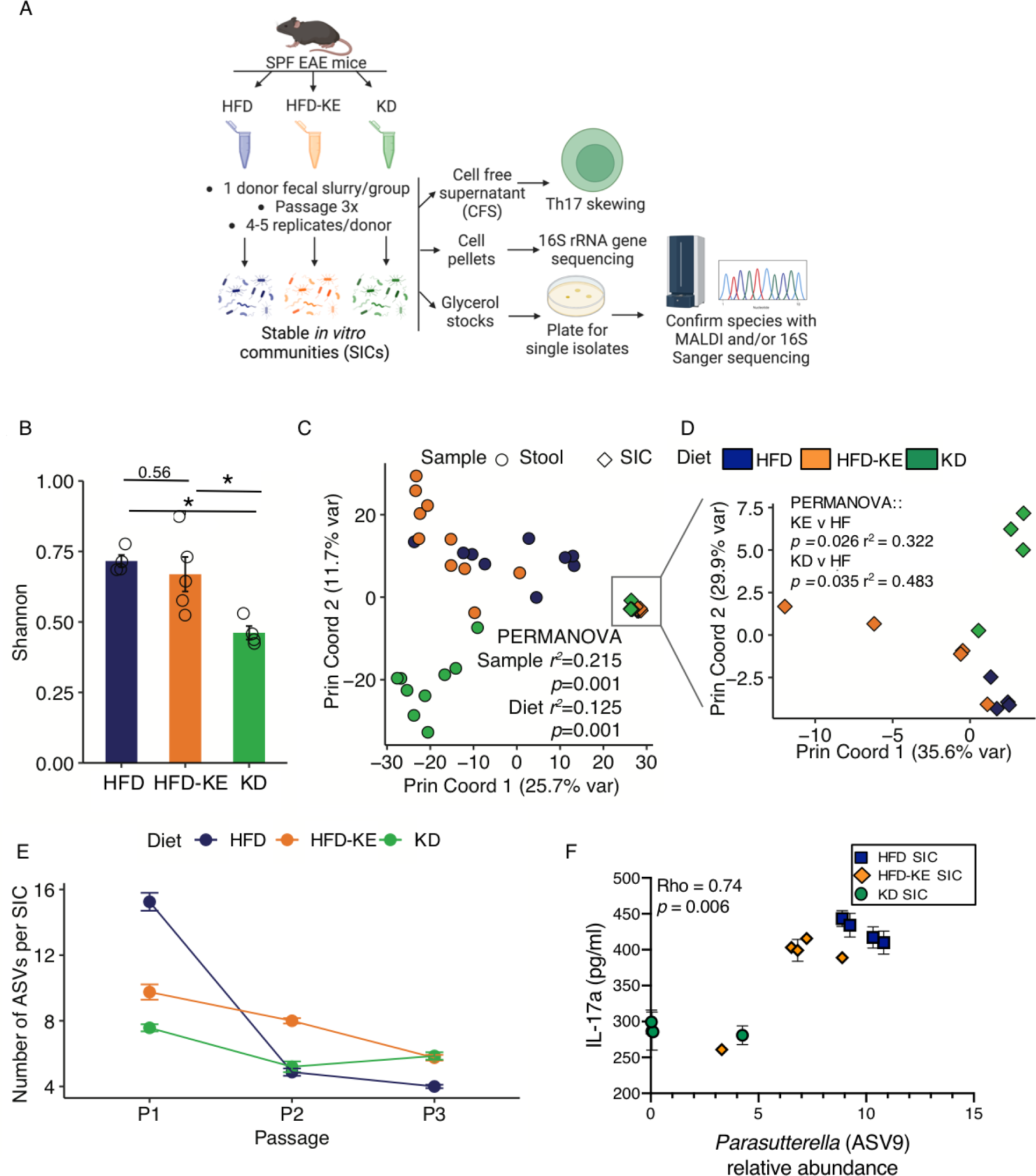
Stable *in vitro* communities maintain community differences based on diet. Related to Figure 5. (A) Stable *in vitro* communities (SICs) were derived from high-fat diet (HFD) or a HFD supplemented with a βHB ketone ester (HFD-KE), or ketogenic diet (KD) mouse stool samples 7 days post EAE induction. Cell free supernatants (CFS) were tested in the Th17 skewing assay. Cell pellets were used for 16S rRNA gene sequencing. We plated from the glycerol stock KD-SIC_4 for single isolates and isolated *L. murinus* strain KD6. (B) Shannon diversity in HFD, HFD-KE, and KD SICs (*p*-values listed; Kruskal-Wallis with Dunn’s test). (C) Principal coordinate analysis (PCA) of Euclidean distances of stool from HFD, HFD-KE, KD fed mice 7 days post-immunization and SICs derived from these samples (*p*-values and r^2^ values listed of sample (stool or SIC) and diet (HFD, HFD-KE, or KD, PERMANOVA). (D) PCA of Euclidean distances of SICs (*p*-values and r^2^ values listed; PERMANOVA). (E) Number of ASVs over passages in HFD, HFD-KE, and KD SICs. (F) ASV9 (annotated as *Parasutterella* by SILVA v138) is positively correlated with IL-17a levels (n=4-5 SICs/group and 3 Th17 skewing replicates; Spearman correlation rho and *p*-values listed; mean±SEM IL-17a levels).

**Figure S7.**
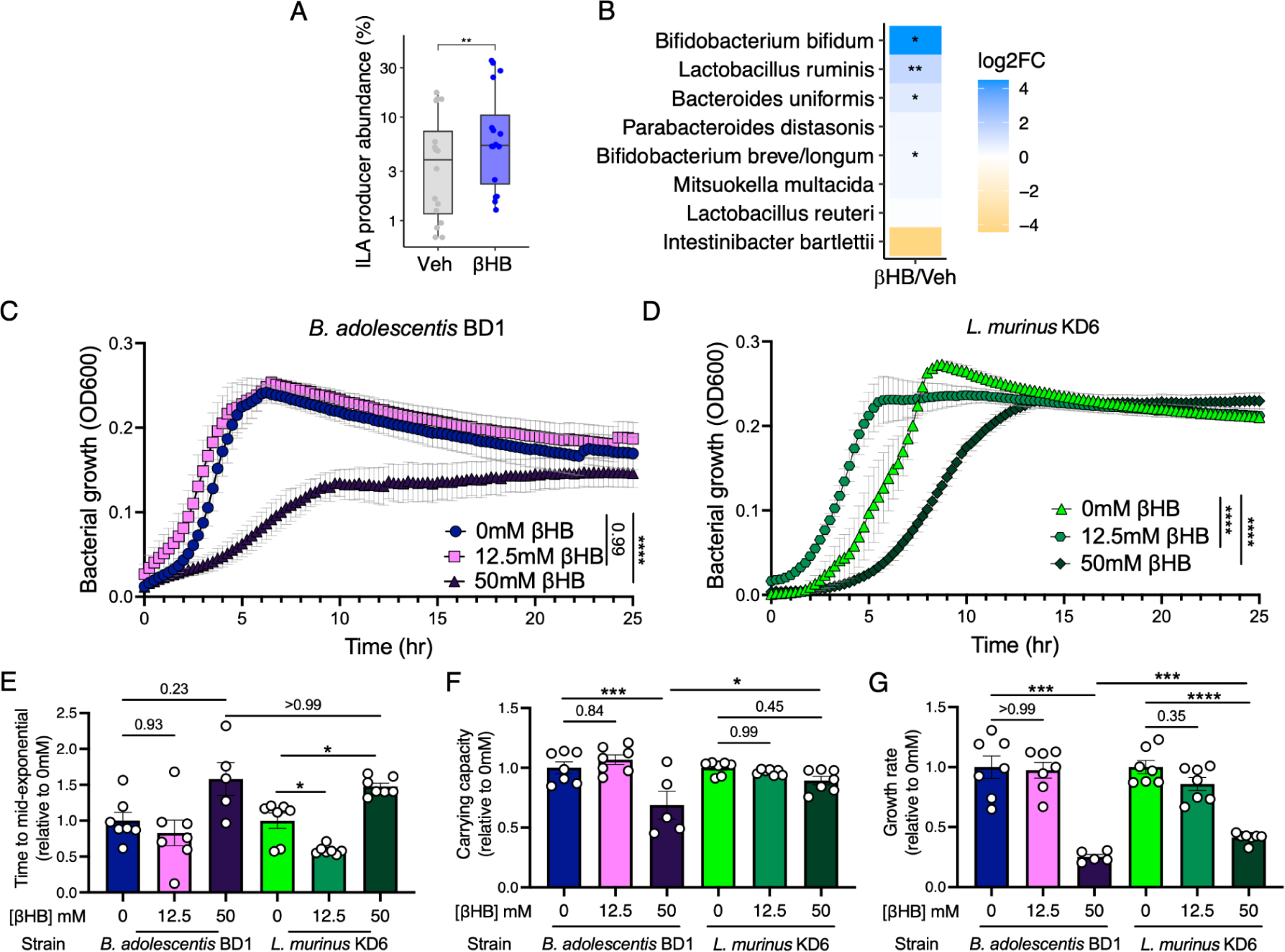
ILA-producing human gut bacteria tolerate βHB. Related to Figure 6. (A) Relative abundance of ILA-producing bacterial species following 48 hours of *ex vivo* incubation of human stool samples with 12.5 mM βHB or vehicle^16^ (n=16 *ex vivo* communities per group; \*\**p*-value<0.01; mixed effects model of abundance, treatment as fixed effect, donor as random effect). (B) Log_2_ fold-change (log_2_ fold-change of βHB/vehicle) of ILA-producing bacterial species in *ex vivo* human stool samples treated with 12.5 mM βHB or vehicle^16^ (n=16 *ex vivo* communities per group; \*\**p*-value<0.01; mixed effects model of abundance, treatment as fixed effect, donor as random effect). (C) *B. adolescentis* BD1 and (D) *L. murinus* KD6 growth in 0mM, 12.5mM, or 25mM βHB (n=6-7; \*\*\*\**p*-value<0.0001; listed; two-way ANOVA; mean±SEM). (E-G) Relative levels of growth metrics compared to the 0mM βHB condition of (E) time to mid-exponential growth, (F) carrying capacity, and (G) growth rate of the curves from **Figure S7C,D** (\**p*-value<0.05; \*\**p*-value<0.01; \*\*\**p*-value<0.0001; \*\*\*\**p*-value<0.0001; listed; Brown-Forsythe ANOVA test; mean±SEM each point represents an individual biological replicate).

